# *CDK13* Mutations Drive Melanoma via Accumulation of Prematurely Terminated Transcripts

**DOI:** 10.1101/824193

**Authors:** Megan L. Insco, Brian J. Abraham, Sara J. Dubbury, Sofia Dust, Constance Wu, Kevin Y. Chen, David Liu, Calvin G. Ludwig, Stanislav Bellaousov, Tania Fabo, Telmo Henriques, Karen Adelman, Matthias Geyer, Phillip A. Sharp, Richard A. Young, Paul L. Boutz, Leonard I. Zon

## Abstract

Transcriptional Cyclin Dependent Kinases modulate RNA Polymerase II function to impact gene expression. Here, we show that *CDK13* is mutated in 4% of patient melanomas and mutation or downregulation is associated with poor overall survival. Mutant CDK13 lacks kinase activity and overexpression in zebrafish leads to accelerated melanoma. CDK13 mutant fish and human melanomas accumulate prematurely terminated RNAs that are translated into truncated proteins. CDK13 binds to and regulates the phosphorylation of ZC3H14, a member of the <u>P</u>oly<u>A</u> e<u>X</u>osome <u>T</u>argeting (PAXT) RNA degradation complex. ZC3H14 phosphorylation recruits the PAXT complex to degrade prematurely terminated polyadenylated transcripts in the nucleus. In the presence of mutant CDK13, ZC3H14 phosphorylation is compromised and consequently fails to recruit the PAXT complex, leading to truncated transcript stabilization. This work establishes a role for CDK13 and the PAXT nuclear RNA degradation complex in cancer and has prognostic significance for melanoma patients with mutated or downregulated CDK13.

---

Transcriptional Cyclin Dependent Kinases (CDKs) are a phylogenetically-related family of kinases that are activated by a cyclin binding partner^1^ and have roles in transcriptional subprocesses including initiation, elongation, and recruitment of complexes engaged in cotranscriptional RNA processing^2–5^. Transcriptional CDKs are being explored as drug targets for many difficult-to-treat cancers^6–9^. Additionally, mutations affecting one of the transcriptional CDKs, CDK13, have recently been found to cause syndromic developmental disorders that affect neural-crest derived tissues^10, 11^ including heart and craniofacial developmental anomalies. However, the role of CDK13 in transcription and RNA processing remains poorly understood and no functional role for CDK13 has been defined in cancer.

RNA polymerase II (RNAPII), the enzyme that transcribes pre-messenger RNA from protein-coding genes, is exquisitely regulated at multiple steps to ensure transcriptional initiation, elongation, and termination are precisely executed. During active elongation, a rapid cascade of a vast number of RNA processing steps takes place^12^. Given this complexity, RNA processing errors can occur, resulting in defective transcripts that must be degraded by the nuclear exosome in order to prevent their translation into truncated proteins^15, 16^.

The nuclear exosome degrades aberrant or unstable nuclear RNAs. Three different human adaptor complexes have been discovered to date, each of which associates with and directs a specific class of nuclear RNAs for exosomal degradation. All three adaptor complexes include the MTR4 helicase, paired with proteins that recognize distinct RNA species. The TRAMP complex targets aberrant nucleolar ribosomal RNAs while the NEXT complex targets PROMoter uPstream Transcripts (PROMPTs) and enhancer RNAs for destruction^13^. The recently reported <u>P</u>oly<u>A</u> tail e<u>X</u>osome <u>T</u>argeting (PAXT) complex degrades prematurely terminated polyadenylated RNAs and requires the adaptor protein ZFC3H1^14, 15^. It is unknown how PAXT-mediated RNA degradation is regulated.

In this study, we show that mutations in *CDK13* occur in melanoma and promote tumor formation and growth. The accelerated tumor progression in melanomas is dependent on Cyclin T1 (CCNT1) rather than the canonical CDK13 cyclin partner, Cyclin K (CCNK). When CDK13 is mutated, ZC3H14 phosphorylation is compromised, and consequently the nuclear RNA exosome fails to be recruited to prematurely terminated polyadenylated RNAs. Failure to recruit the nuclear RNA exosome to prematurely truncated transcripts results in accumulation of aberrant RNAs and the production of truncated proteins in melanoma. This work uncovers a role for the PAXT nuclear RNA exosome in human cancer and defines its regulation by CDK13.

## Results

### CDK13 is a melanoma tumor suppressor

We examined the transcriptional CDK loci in melanoma patient samples and found *CDK13* is mutated in 4% (22/520 cases) of cutaneous melanomas, with 9 distinct missense mutations occurring in the kinase domain, and 7 of these mutations near the ATP-binding site (Figure 1A). Surprisingly, many of the same amino acid changes were found in CDK13-associated developmental syndromes^10^, suggesting a specific activity of these mutations. Survival analysis of human TCGA data revealed that *CDK13* downregulation (*z* score ≤ −1.0) or mutation (*CDK13-*altered) correlated with decreased overall survival [median *CDK13-*altered survival of 48.8 months vs. remaining case survival of 102 months ^16, 17^ (Figure 1B)]. Patients initially staged with 0/1/2 melanoma that had *CDK13*-alterations exhibited reduced overall survival compared to remaining stage 1/2 melanoma patients (Figure S1A). *CDK13* melanoma mutations near the ATP-binding site were plotted onto the crystal structure (Figure 1C)^18^ and 6/7 of these mutations were predicted to eliminate kinase activity by disruption of T-loop activation (R860Q or P869S), alteration of the substrate binding pocket (W878L), or disruption of the kinase domain structure (P881L, P893L, and I843N). *In vitro*, CDK13^WT^ activated by the typical cyclin partner, CCNK, had robust kinase activity while the R860Q, W878L, and K734R mutants failed to phosphorylate full-length RNAPII C-terminal domain (CTD) and a second substrate (Figure 1D, S1B). As a control, we included the catalytically-dead CDK13^K734R^ mutation, which replaces the lysine required for catalysis of all known kinases. In summary, these data suggest that *CDK13* is mutated in melanoma, mutations abrogate CDK13 kinase activity, and mutation or downregulation of *CDK13* in patient melanomas is associated with decreased overall survival.

**Figure 1:**
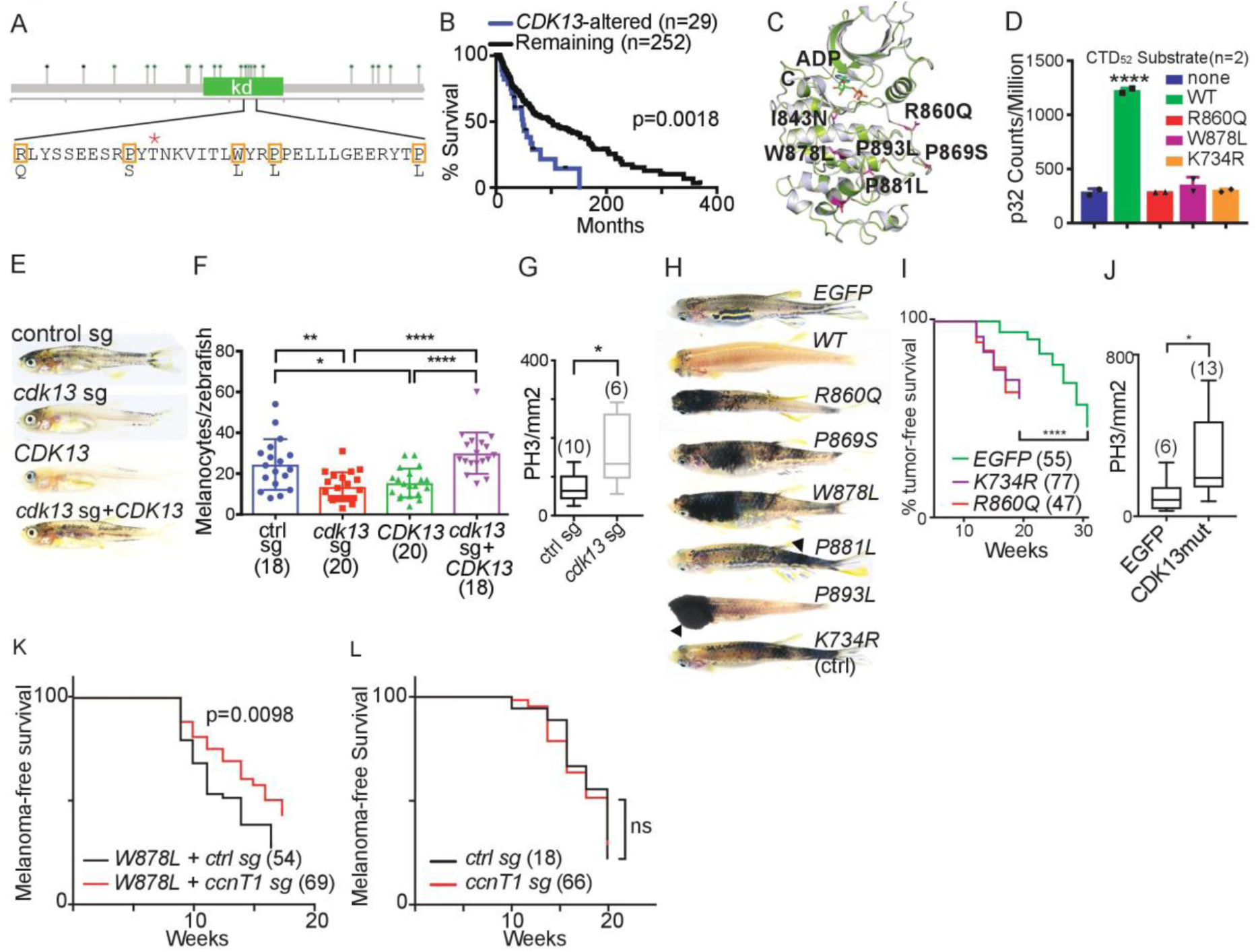
CDK13 is a melanoma tumor suppressor. A) *CDK13* mutation plot in melanoma. Orange rectangles denote 5/7 mutations near the ATP-binding site. *=phosphorylation site. B) Survival plot of *CDK13* downregulation or mutation in patients. p=0.0018 Log-rank. n= patients. C) Patient kinase-domain mutations mapped on the CDK13 crystal structure. Red residues = mutations. D) *In vitro* kinase assay of wild type and patient-mutated CDK13 using full-length CTD52 as the substrate. One-way ANOVA with no kinase vs. all conditions; WT CDK13 ****=q=0.0001, all mutated CDK13 comparisons non-significant. Mean +SD. n=2 replicates. E) Representative 4 week photos of *mitfa:BRAF; p53-/-; mitfa-/-* zebrafish injected with 1) control gRNA, 2) *cdk13* gRNA, 3) overexpression of human *WT CDK13*, or *cdk13* gRNA and overexpression of wild-type *hCDK13*. F) Quantification of melanocytes at 3 days post fertilization for *mitfa:BRAF; p53-/-; mitfa-/-* zebrafish injected with 1) control CRISPR (blue), 2) *cdk13* gRNA (red), 3) overexpression of human *CDK13^WT^* (green), or *cdk13* gRNA and overexpression of *CDK13^WT^* (purple). p=one-way ANOVA, multiple comparisons. Mean +/-SD. *= q=0.186, **=q=0.0030, ****=q<0.0001. (#)=zebrafish. G) Phospho Histone H3 Serine 10 (PH3) antibody staining/mm^2^ of melanomas from *mitfa:BRAF; p53-/-; mitfa-/-* zebrafish with *cdk13* gRNA compared with control gRNA. p=0.014 (Mann Whitney two-tailed t test). (#)=melanomas. Box and whiskers plot, min to max. H) 9-week photos from melanocyte-specific expression of 1) EGFP, 2) wild-type human *CDK13*, 3-7) *CDK13* patient mutations, and 8) *CDK13 K734R* (catalytically dead) in *BRAF; p53-/-*; *mitfa-/-* zebrafish. Arrows=melanomas. I) % melanoma-free survival of *BRAF; p53-/-; mitfa-/-* zebrafish injected with melanocyte-specific expression of *EGFP, CDK13^R860Q^ (*patient mutation), and *CDK13^K734R^ (*catalytically dead). ****=p<0.0001 (log-rank). (#)=zebrafish. J) PH3 antibody staining/mm^2^ of melanoma from *mitfa:BRAF; p53-/-; mitfa-/-* zebrafish with *EGFP* vs. *CDK13* mutant (*W878L* or *P893L)*. p=0.0125 (Mann Whitney two-tailed t test). (#)=melanomas. Box and whiskers plot, min to max. K) Melanoma-free survival of melanocyte-specific expression of *CDK13 W878L* coinjected with melanocyte-specific CRISPR of either a control gRNA or *ccnT1* gRNA. p=0.0098 (log-rank). (#)=zebrafish. L) Melanoma-free survival of melanocyte-specific expression of control gRNA or *ccnT1* gRNA alone. ns=non significant (log-rank). (#)=zebrafish. See also Figure S1.

To examine *cdk13* loss in melanoma, we used a *BRAF^V600E^; p53-/-; mitfa-/-* zebrafish melanoma model^19^ in which Cas9 was expressed in melanocytes using the *mitfa* promoter and either a *cdk13* gRNA or a control gRNA was ubiquitously expressed. Zebrafish lacking *cdk13* showed significantly decreased numbers of melanocytes as compared with zebrafish expressing a control gRNA, indicating that *cdk13* is required for normal melanocyte development (Figure 1E, 1F, S1D vs. S1C). Overexpression of human wild-type *CDK13* (*CDK13^WT^*) resulted in fewer melanocytes during development (Figure 1E, 1F, S1E). Injection of a *cdk13* gRNA/Cas9-expressing vector and human *CDK13^WT^* rescued melanocyte numbers in the melanoma model (Figure 1E, 1F, S1F). These data show that tightly-controlled levels of *cdk13* are required for melanocyte development and that human *CDK13* can complement the loss of the zebrafish gene.

Rarely, melanomas arose in zebrafish with *cdk13* melanocyte-specific CRISPR deletion, and these tumors appeared to grow faster than melanomas with intact *cdk13*. Phospho-histone 3 (PH3) immunohistochemical (IHC) staining of melanomas with *cdk13* gRNA and a control gRNA revealed that *cdk13* CRISPR-deleted tumors were significantly more proliferative than melanomas with retained *cdk13* (Figure 1G, S1H). PCR across the CRISPR cut site confirmed frameshift indels, and more in-frame indels were observed in *cdk13* gRNA compared to control gRNA melanomas (Figure S1G). These data suggest that melanocytes with loss of *cdk13* may be selected against during development, but surviving melanocytes give rise to highly proliferative melanomas.

To test whether CDK13 melanoma-associated mutations cause more aggressive melanoma *in vivo*, mutant CDK13 was expressed in melanocytes in a *BRAFV600E; p53-/-; mitfa-/-* zebrafish melanoma model^20, 21^. Control melanocyte-specific *EGFP* expression resulted in expected mosaic zebrafish stripes, whereas human CDK13^WT^ expression resulted in dramatically fewer melanocytes at 9wpf (Figure 1H, S1I). In contrast, melanocyte-specific overexpression of CDK13^R860Q^, CDK13^P869S^, CDK13^W878L^, CDK13^P881L^, and CDK13^P893L^ caused the appearance of black patches at 9wpf (Figure 1H, S1I) and expedited tumor onset (Figure 1I, S1J). Overexpression of the catalytically-dead CDK13^K734R^ mutant also caused black patches at 9wpf and expedited tumor onset (Figure 1H, 1I, S1I). CDK13 W878L-and P893L-expressing melanomas had more PH3-positive cells than control EGFP melanomas by IHC (Figure 1J, S1K). Because all *CDK13* mutations promoted melanoma to a similar degree, we have used them interchangeably in most assays, and refer to them collectively hereafter as *CDK13^mel^*. Since both CRISPR-mediated deletion and CDK13^mel^ expression in zebrafish caused more proliferative melanomas, our data suggest that CDK13^mel^ mutations act through a dominant negative mechanism.

As CDK13^mel^ acts via a dominant negative mechanism and no *CDK13^mel^* mutations are observed in the Cyclin binding domain, we hypothesized that Cyclin binding is required for CDK13^mel^ dominant negative activity. To test whether the canonical CDK13 Cyclin, CCNK^18, 22, 23^, or the related CCNT1 are required for CDK13^mel^ dominant negative melanomagenesis *in vivo*, we co-injected vectors that express melanocyte-specific CDK13^mel^ and melanocyte-specific CRISPR of *ccnK* or *ccnT1* in the *mitfa^-/-^; BRAF; p53^-/-^* zebrafish melanoma model. *ccnK* melanocyte-specific CRISPR expedited melanoma in the presence of CDK13^W878L^, but not alone (Figure S1L-M). *ccnT1* melanocyte-specific CRISPR suppressed CDK13^W878L^ and CDK13^R860Q^ oncogenesis but had no effect by itself (Figure 1K-L, S1O). PCR across the CRISPR cut site confirmed indels directed by the *ccnT1, ccnK*, and control gRNAs (Figure S1N). These data indicate that CDK13^mel^ oncogenesis requires CCNT1 for its dominant negative oncogenic activity.

### Mutant CDK13 binds chromatin in a dominant negative manner

As CDK13 is localized in the nucleus^24^ and CDK13^WT^ can phosphorylate the RNAPII CTD (Figure 1D), chromatin immunoprecipitation sequencing (ChIP-seq) was used to determine the localization of CDK13^mel^ on melanoma chromatin. ChIP-seq was performed on approximately equal cell numbers gathered from EGFP and CDK13^W878L^*-*expressing zebrafish melanomas (Figure 2A). EGFP*-*expressing melanomas were used as the ChIP-seq control since zebrafish expressing CDK13^WT^ had very few melanocytes and rarely developed melanoma. ChIP-seq from EGFP melanomas showed zebrafish Cdk13 was especially enriched at transcriptional start sites (TSSs) (Figure 2B, left, green). CDK13^W878L^ overexpression in melanoma resulted in increased CDK13 signal at TSSs (Figure 2B, left, black). CDK13^W878L^ bound to genes with Cdk13^WT^ occupancy in control melanomas and the amount of CDK13^W878L^ bound correlated with the amount of Cdk13^WT^ bound in control melanomas (Figure 2C).

**Figure 2:**
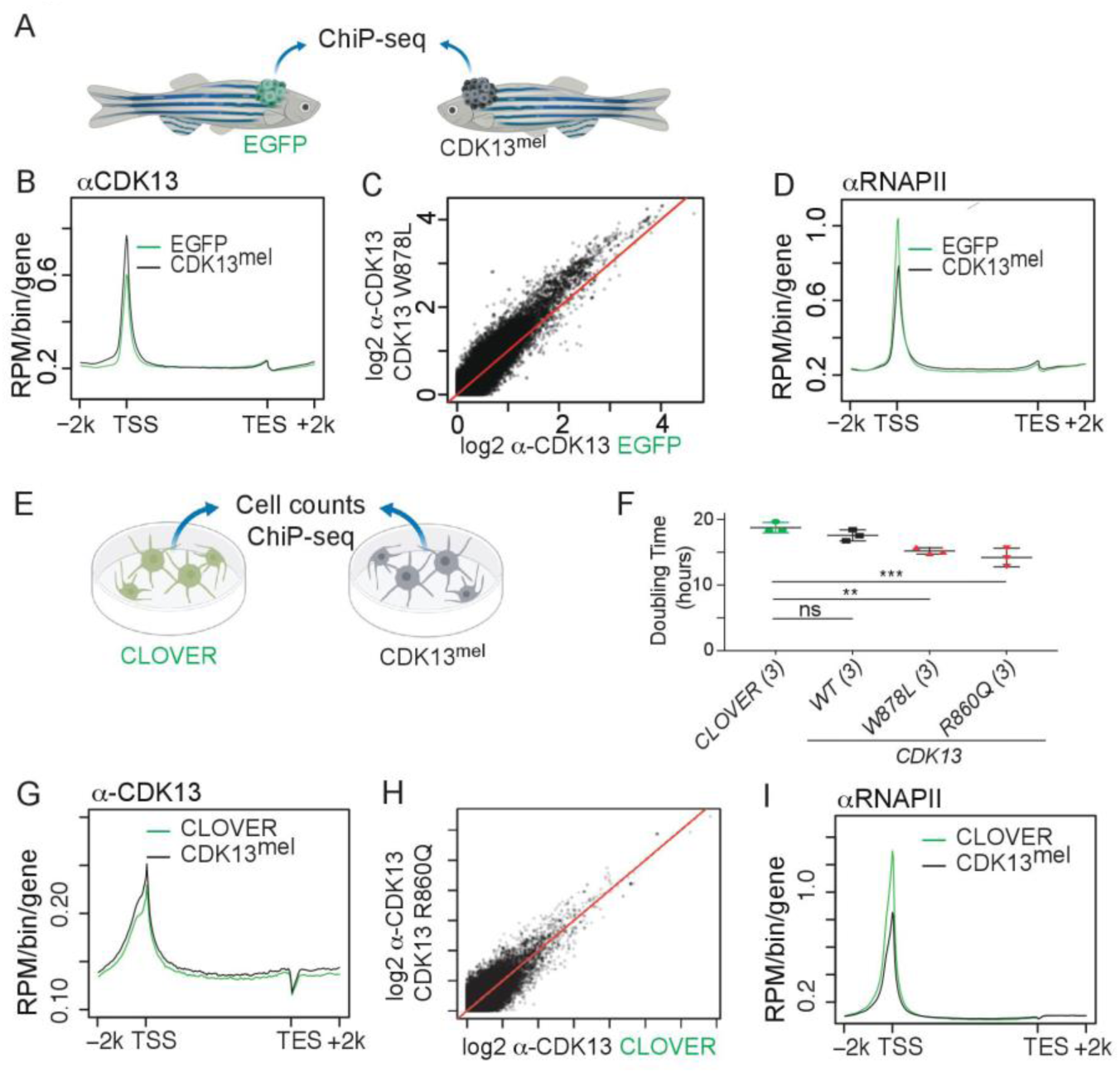
Mutant CDK13 binds chromatin in a dominant negative manner. A) Schematic for zebrafish melanoma chromatin immunoprecipitation sequencing (ChIP-seq) of *mitfa:BRAF; p53-/-; mitfa-/-* melanomas with melanocyte-specific expression of *EGFP* (green) or *CDK13^W878L^* (black). B-C) anti-CDK13 zebrafish melanoma ChIP-seq displayed as a B) metagene or C) log2 from *EGFP* vs. *CDK13^W878L^* for each gene. D) anti-RNAPII zebrafish melanoma ChIP-seq metagene. E) Schematic for ChIP-seq from human melanoma cells expressing CLOVER fluorescent protein or CDK13^mel^. F) Doubling time for human melanoma cell lines expressing CLOVER, CDK13^WT^, CDK13^W878L^, or CDK13^R860Q^. One-way ANOVA with CLOVER vs. all conditions. CDK13^W878L^ q=0.0044, CDK13^R860Q^ q=0.0009. Others non-significant. Mean +SD. n=3 biologic replicates. G-H) anti-CDK13 human melanoma cell ChIP-seq displayed as a G) metagene or H) log2 from *EGFP* vs. *CDK13^W878L^* for each gene. I) anti-RNAPII human melanoma cell ChIP-seq metagene. See also Figure S2.

To determine whether mutant CDK13 also bound chromatin in human melanoma cells, CLOVER fluorescent protein, CDK13^WT^, CDK13^W878L^, or CDK13^R860Q^ were expressed in human A375 melanoma cells (Figure 2E). Cells expressing either CDK13^mel^ protein grew more quickly than cells expressing CDK13^WT^ or CLOVER (Figure 2F, Figure S2A). CDK13^R860Q^ also bound chromatin loci previously bound by CDK13^WT^, similar to the findings in zebrafish melanomas (Figure 2G-H). Because CDK13^mel^ binds chromatin at sites normally bound by CDK13^WT^, CDK13^mel^ is positioned to interfere with CDK13^WT^ kinase activity and thus act in a dominant negative manner in zebrafish and human melanoma cells.

As CDK13^mel^ bound chromatin, we next sought to determine if CDK13^mel^ disrupted RNAPII occupancy. ChIP-seq was performed with antibodies that recognize either RNAPII (anti-8WG16), or RNAPII phosphorylated at the CTD Serine-2 position (anti-Ser2p), a modification of elongating RNAPII involved in the recruitment of RNA processing machinery ^12, 25, 26^. Zebrafish melanomas expressing CDK13^W878L^ exhibited perturbed RNAPII as compared to control melanomas, with a notable decrease in TSS-proximal RNAPII and a slight increase gene body RNAPII (Figure 2D). Ser2p RNAPII occupancy was also perturbed; genes with the highest increase in CDK13^mel^ binding had increased Ser2p (Figure S2B, left). When the Ser2p RNAPII signal was normalized to total RNAPII, a dramatic increase in Ser2P RNAPII was observed at the promotor-proximal region (Figure S2C, left). Promoter-proximal enrichment of Ser2P RNAPII has been reported for conditions causing slowed RNAPII^27^. Human melanoma cell CDK13^R860Q^ expression caused similar perturbations in RNAPII occupancy [Figure 2I, S2B (right), S2C (right)]. To ascertain whether CDK13^mel^ expression impacts transcription based on gene expression level or gene length, we compared the RNAPII and Ser2P ChIP signals in gene subsets partitioned by gene expression level or gene length. More CDK13^mel^ was bound and RNAPII was more perturbed on highly expressed genes (Figure S2D-E), however there was no correlation with gene length (S2F-G). These data show CDK13^mel^ expression perturbed RNAPII occupancy on highly expressed genes, suggesting that CDK13 normally acts via a mechanism important on highly expressed genes.

Notably, changes in the distribution of the Ser2p signal upon CDK13^mel^ expression differed distinctly from those observed upon loss of a close paralog, CDK12. In cells lacking CDK12, Ser2p was strongly decreased across the gene body despite an increase in total RNAPII^28^. These data also suggest that CDK13^mel^ promotes oncogenesis via a mechanism that is distinct from CDK12.

### Mutant CDK13 causes accumulation of prematurely-terminated RNAs

To determine the transcriptional impact of CDK13^mel^ in zebrafish melanomas, RNA sequencing (RNA-seq) was conducted on *BRAF; p53-/-; mitfa-/-* zebrafish melanomas expressing *EGFP* (n=4) or mutant *CDK13* (*R860Q*, n=5; *K734R*, n=5, Figure S3A). Differential expression reassuringly showed the most differentially expressed gene in the zebrafish melanomas was human *CDK13^mel^*. 158 downregulated and 53 upregulated genes with a *q value* <0.05 were identified. Panther GO-Slim Biologic Process pathway analysis of upregulated genes showed no pathway enrichment, while downregulated genes were enriched for 1) positive regulation of GTPase activity (q=1.63×10-7), 2) cellular response to tumor necrosis factor (TNF) (q=1.64×10-7), and 3) granulocyte chemotaxis (q=1.61×10-7).

As CDK13 is phylogenetically related to known transcriptional kinases CDK9 and CDK12, and CDK13^mel^ expression caused a perturbation in RNAPII occupancy, we tested for a global RNA defect by quantifying differential exon expression. RNA differential expression analysis of first (F), alternative first (AF), internal (I), last (L), or alternative last (AL) exons in each gene were quantified genome-wide in zebrafish melanomas. Many genes exhibited increased read coverage in the first exon as compared to the last exon in CDK13^mel^ expressing as compared with control melanomas (p=2.2×10^-16^, two-sided Wilcoxon rank sum test) (Figure 3A). These data show that prematurely terminated transcripts containing the 5’ but not the 3’ end of the gene are present in CDK13^mel^ zebrafish melanomas.

**Figure 3:**
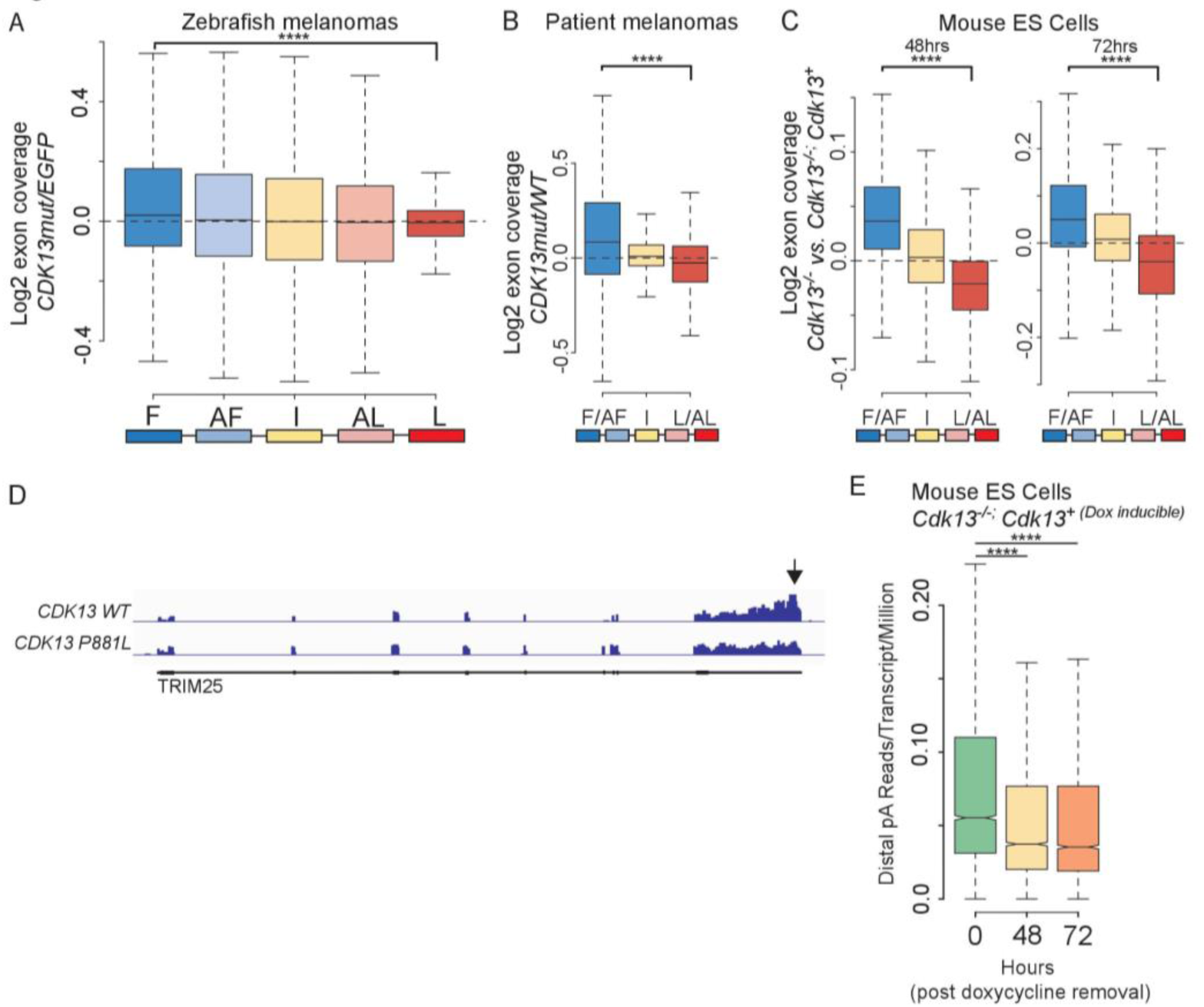
CDK13 mutation/depletion results in accumulation of prematurely terminated RNAs. A) Distributions of the log2-fold differences in normalized exon coverage for each exon category in *CDK13 R860Q* (n=5) vs. EGFP (n=4) expressing zebrafish melanomas. F=first exon. AF=alternative first exon. I=internal exon. AL=alternative last exon. L=last exon. ****=p<2.2×10-16, F vs. L exon. B) As in (A) plotted for *CDK13* mutant (n=3) vs. *CDK13 WT* (n=5) matched control patient melanomas. ****=p<2.2×10-16, F/AF vs. L/AL exon. C) As in (A) plotted for *Cdk13-/-* (n=4) vs. *Cdk13-/-; Cdk13+* mouse embryonic stem (ES) cells at 48 hours (left, n=4) and 72 hours (right, n=4). ****=p<2.2×10-16, F/AF vs. L/AL exon. D) Example IGV track of melanoma patient RNA-seq for target gene TRIM25 from a patient with a *CDK13 ^P881L^* mutation and a matched patient with CDK13^WT^. Arrow highlights 3’ exon coverage difference. E) Distribution of the ratios of reads representing distal (full-length) polyadenylation site usage for each gene per transcript of that gene present in every million transcripts in the sample (polyA reads/transcripts per million (TPM)) in *Cdk13-/-; Cdk13+* mouse ES cells 0, 48, and 72 hours after Cdk13^+^ depletion. ****=p<2.2 x10-16. In all box plots, the black horizontal line indicates the median and whiskers extend to 1.5 x the interquartile range. p values were determined by two sided Wilcoxon rank sum test. See also Figure S3.

To test whether *CDK13^mel^* mutations have a similar effect on RNAs in patient melanomas, we analyzed polyA-selected RNAseq data from TCGA data. Melanomas with kinase domain mutations (n=3) were compared to matched wild-type controls (n=5). Similar to zebrafish melanomas, increased read coverage in the first exon versus the last exon was observed in *CDK13^mel^* as compared to *CDK13^WT^* patients (p=2.2×10^-16^, two-sided Wilcoxon rank sum test) (Figure 3B). One example gene that was significantly affected in patient samples was TRIM25 (Figure 3D). The evidence for prematurely terminated RNAs was of greater magnitude in polyA-selected RNA-seq, consistent with the prematurely terminated transcripts being polyadenylated. These data demonstrate that CDK13^mel^ expression in both zebrafish and human melanomas results in the accumulation of prematurely terminated polyadenylated transcripts.

To determine whether genetic deletion of *CDK13* also results in accumulation of prematurely terminated transcripts, we generated mouse embryonic stem cells (mESCs) that were *Cdk13-/-* and carried a doxycycline-inducible *Cdk13* rescue transgene. After removal of doxycycline for 0,48, or 72 hours, polyA-selected RNA was subjected to stranded sequencing (Figure S3B-E). Similar to zebrafish and patient melanomas expressing CDK13^mel^, increased first exon coverage and decreased last exon coverage was observed at 48 hours as compared to 0 hours in Cdk13-depleted mESCs. Cdk13-depleted mESCs exhibited an additional increase in the ratio of first to last exon coverage after 72 hours (Figure 3C). This data further supports the hypothesis that Cdk13 is required to prevent accumulation of prematurely terminated polyadenylated transcripts.

Notably, in contrast to the increase in distinctly positioned intronic polyadenylation events observed upon Cdk12 depletion^28^, we were unable to detect cleavage and polyadenylation at specific sites within gene bodies. We developed an algorithm to quantify sequencing reads with non-genomically encoded polyA tails that map to the distal polyA sites of genes. Using this algorithm, we observed a significant decrease in the ratio of transcripts from highly expressed polyadenylated genes at the distal polyA site (corresponding to the full-length mRNA) to total transcript number in Cdk13-depleted mESCs (Figure 3E). Cdk13 depletion causes RNAs terminating upstream of the normal polyA site to be either generated more frequently or degraded more slowly. As a consequence, the pool of cellular transcripts contains proportionately fewer full-length mRNAs.

### Mutant *CDK13* causes accumulation of truncated proteins

Prematurely terminated transcripts are typically degraded by the nuclear exosome, which prevents them from being exported and translated. To determine whether the prematurely terminated transcripts we observed in CDK13^mel^ zebrafish melanomas are translated, we used tandem mass spectrometry to analyze global protein expression in melanomas isolated from age-matched *BRAF^V600E^; p53-/-; mitfa-/-* zebrafish with melanocyte-specific expression of CDK13^mel^ (n=3) or EGFP (n=3). To identify consistent changes across replicates, peptides measurements with a t-test result of p<0.1 were mapped. The log2-fold change in peptide measurements between CDK13^mel^/EGFP were binned and plotted along protein length to create a metaplot (Figure 4A, r= −0.5349, p value=0.0001). The negative slope in the metaplot is consistent with an increase in short proteins arising from the translation of prematurely terminated polyadenylated transcripts. These data suggest that rather than being degraded by the nuclear exosome, prematurely terminated transcripts accumulate and are translated into truncated proteins in CDK13^mel^ expressing cells.

**Figure 4:**
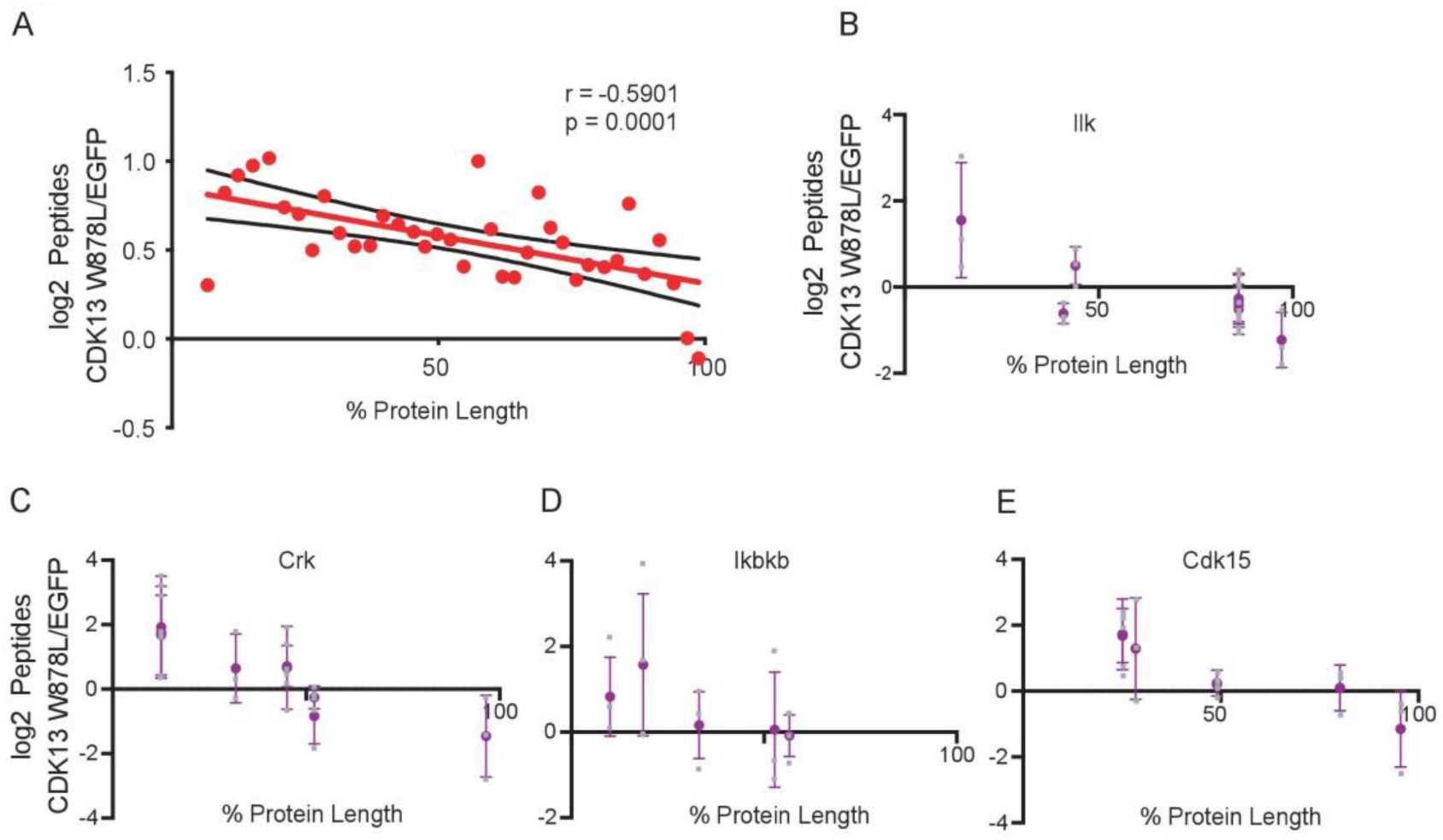
Mutant *CDK13* caused accumulation of truncated proteins. A) Log2 peptide measurement ratio between cells bearing CDK13^mel^ and EGFP plotted across a normalized protein length. r = −0.5901, p = 0.001. n = 3 primary zebrafish melanomas from each condition. B-E) Log2 *CDK13^mel^/EGFP* zebrafish melanoma tandem mass spectrometry peptide measurements plotted by % protein length for candidate dominant negative protein targets B) Ilk, C) Crk, D) Ikbkb, and E) Cdk15. Gray squares = individual measurements. Error bars = SD. n=3 zebrafish melanomas of each genotype. See also Figure S4.

To identify specific proteins that are most impacted by CDK13^mel^, two methods were used. First, we assessed proteins as shown in Figure 4A that had >1 measurement and metaplot slope <-1, which identified 263 truncated proteins. Second, using all data, proteins with >3 peptide measurements were tested for a negative slope (log^2^ CDK13^mel^/EGFP, F-test p<0.05). This more restrictive analysis identified 103 candidates (Figure S4), 56 of which were distinct from the first group. Of these 103 candidate proteins, the most significantly enriched KEGG Pathways were metabolic ribosome (FDR 2.01e-07) and carbon metabolism (3.01 e-05). Several of the truncated proteins we identified are predicted to lose carboxy-terminal enzymatic domains, including Ilk, Crk, Ikbkb, and Cdk15 (Figure 4B-E). Ilk, Crk, and Ikbkb have published evidence of dominant negative activity upon carboxy-terminal loss^29–31^. Individual impacted proteins showed an increase in truncated species at the expense of full-length proteins in CDK13^mel^ zebrafish melanomas. These data indicate that CDK13^mel^ expressing zebrafish melanomas translate prematurely terminated transcripts to produce truncated proteins, which we hypothesize cause CDK13^mel^ oncogenesis via tumor suppressor loss and/or expression of dominant negative protein truncations.

To determine whether truncated protein production correlated with CDK13^mel^ binding at the gene locus, we overlaid the 319 truncated proteins with the top quintile of genes with enriched CDK13^mel^ binding as determined by ChIP-seq. 133/319 truncated proteins (42%) were within the top quintile of CDK13^mel^ enrichment. As the ChIP-seq and tandem mass spectrometry experiments were done on different cohorts of primary zebrafish melanomas, and given the detection limits of mass spectrometry, the overlap in this restrictive analysis suggests that genes with high levels of CDK13^mel^ chromatin binding are more likely to accumulate prematurely terminated RNAs which are translated into truncated proteins than would be expected by chance (p<0.0001, Fisher’s exact test).

### Mutant CDK13 disrupts the polyA RNA exosome

To elucidate how CDK13^mel^ expression results in the accumulation of truncated RNAs in human melanoma cells, V5-tagged CDK13^WT^ (n=3), CDK13^W878L^ (n=2), or control fluorescent protein CLOVER (n=3) were immunoprecipitated from nuclear extracts and co-immunoprecipitating proteins were identified by mass spectrometry (IP-MS) (Figure 5A). In the CDK13^WT^ IP, of the 11 co-precipitating proteins identified (Figure 5C), 9 are RNA binding proteins (*q*=3.04×10^-5^) and 3 are involved in negative regulation of polyadenylated RNA (*q*=2.73×10^-4^) ^32, 33^. We found both CDK13^WT^ and CDK13^W878L^ bound to CCNT1, again implicating CCNT1 as the required cyclin-binding partners for CDK13 in human melanoma cells as our functional data suggested (Figure 2J-L); surprisingly, the canonical cyclin binding partner of CDK13, CCNK, was not detected above background^18, 22, 34^. This unbiased approach indicated that the most significant interacting partners of CDK13 are CCNT1 and multiple factors involved in RNA processing.

**Figure 5:**
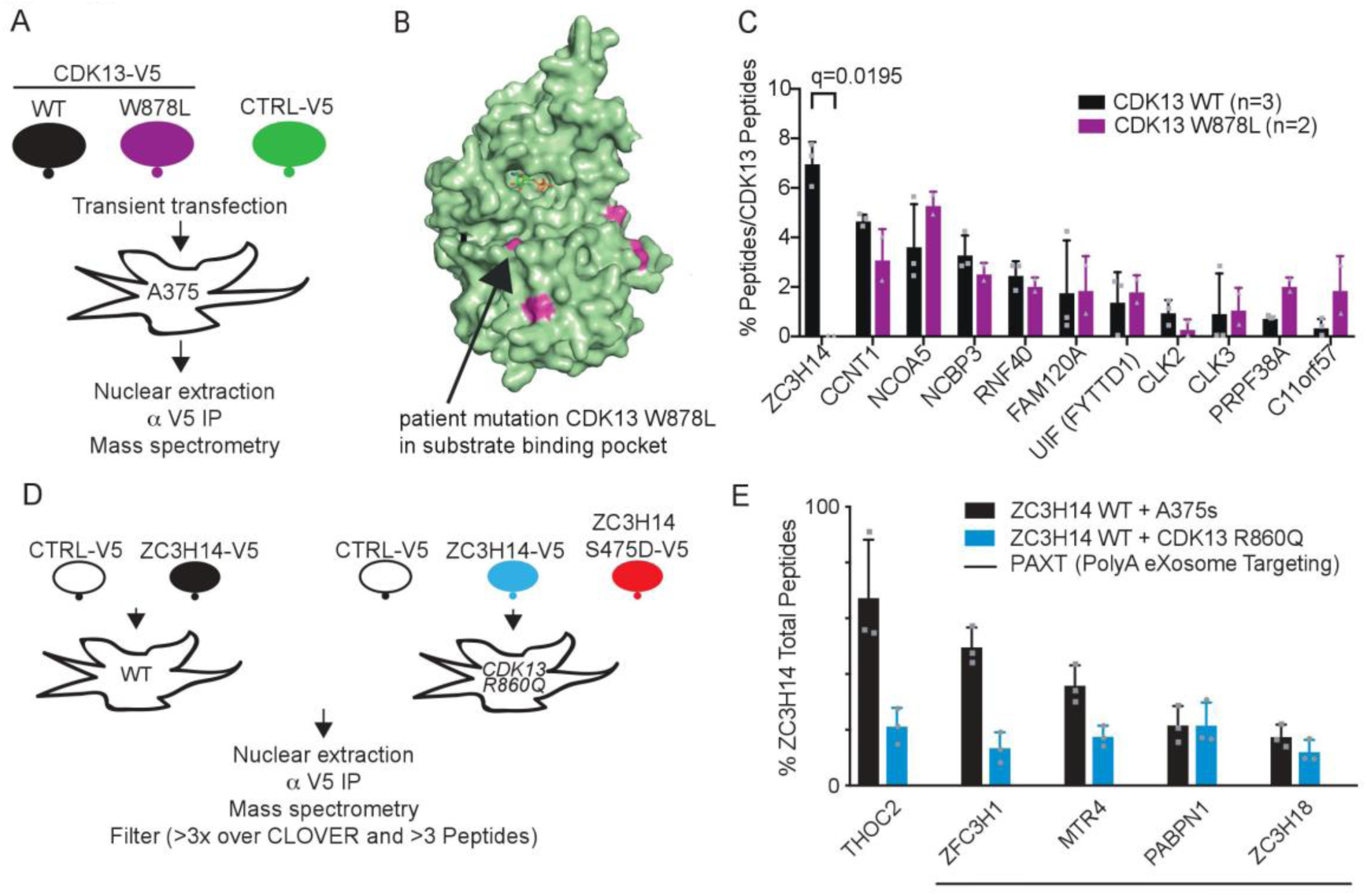
Mutant CDK13 disrupts the polyA RNA exosome. A) Immunoprecipitation mass spectrometry (IP-MS) scheme. B) Surface display of *W878L* patient mutation in substrate binding groove. C) IP-MS most abundant proteins isolated as a percent of CDK13 peptides isolated. q=unpaired t test with multiple test correction. Mean +SD. n=biologic replicates. Gray dots = individual values. D) ZC3H14 IP-MS scheme in CDK13 mutant expressing human melanoma cells. E) IP-MS of PAXT complex proteins isolated as a percent of ZC3H14 peptides isolated. Mean +SD. n=biologic replicates. Gray dots = individual values. See also Figure S5.

In the CDK13^WT^ IP, the most abundantly identified interacting protein was ZC3H14, a zinc finger protein implicated in negative regulation of polyA tail length^35, 36^. ZC3H14 was also recently identified as part of the <u>P</u>oly<u>A</u> tail e<u>X</u>osome <u>T</u>argeting (PAXT) complex^14^ which is responsible for targeting prematurely terminated polyadenylated transcripts to the nuclear exosome for degradation^15^. Of the mutations observed in melanoma patients, *CDK13^W878L^* was chosen because it changes an amino acid in the substrate binding groove, which we hypothesized would disrupt substrate binding (Figure 5B). In the CDK13^W878L^ IP, the ZC3H14 interaction was strongly disrupted (*q*=0.0195) while all other protein interactors were unaffected (Figure 5C). These data suggest that ZC3H14 binds CDK13 in the substrate binding groove and could be a CDK13 substrate.

To identify ZC3H14 binding partners in CDK13^mel^ cells, the nuclear isoform of ZC3H14 and a control protein were tagged and transiently expressed in CDK13^mel^ and CDK13^WT^ human melanoma cells (Figure 5D). By IP-MS, we found that the three most abundant binding partners for ZC3H14 in CDK13^WT^ cells were THOC2, ZFC3H1, and MTR4 (Figure 5E, black bars). THOC2 functions in RNA export, ZFC3H1 functions as a linker between the PAXT and the nuclear RNA degradation complex, and MTR4 is a helicase required for the function of all three known nuclear RNA degradation complexes^14^. Other members of the PAXT complex were identified including PABPN1 and ZC3H18. In cells expressing CDK13^mel^, ZC3H14 lost phosphorylation at S475 (as determined by mass spectrometry) (Figure S5A) and exhibited decreased binding to THOC2 (q<0.0001), ZFC3H1 (q<0.0001), and MTR4 (q=0.018) but stable binding to PABPN1 and ZC3H18 (Figure 5E, blue bars). Together, these data show that CDK13 normally activates the recruitment of the PAXT complex to prematurely terminated polyadenylated transcripts via phosphorylation of the ZC3H14 adaptor. Upon loss of CDK13 function, the PAXT complex is not efficiently recruited to truncated mRNAs, resulting in their stabilization and translation into truncated polypeptides that promote melanomagenesis (Figure S5B).

## Discussion

Our data show that CDK13 functions as a tumor suppressor in melanoma and that CDK13^mel^ alleles promote melanomagenesis through a dominant negative mechanism requiring CCNT1. We provide evidence that *CDK13* mutation or loss leads to a failure to degrade prematurely terminated transcripts. We found that CDK13 binds to, and may phosphorylate ZC3H14, a component of the PAXT complex which is responsible for degrading polyadenylated prematurely terminated RNAs. Consistent with the functional link to CDK13 that we have established, ZC3H14 germline mutations also cause a neurodevelopmental disorder^37^. We confirmed that ZC3H14 associates with the PAXT nuclear RNA exosome complex as recently reported^14^. Upon CDK13^mel^ expression, we showed that ZC3H14 lost phosphorylation at S475, which coincided with reduced recruitment of ZFC3H1 and MTR4. Because PAXT has been shown to specifically target polyadenylated prematurely terminated RNAs for nuclear degradation^15^, we suppose that CDK13 normally phosphorylates ZC3H14 to recruit the PAXT complex to degrade aberrant RNAs made during failed transcriptional elongation. When *CDK13* is mutated, failure to recruit the PAXT complex results in the accumulation of truncated transcripts which are in turn translated into truncated proteins, contributing to aggressive melanoma (Figure S5B).

The proteomic data suggest prematurely terminated transcripts that accumulate in CDK13^mel^ expressing cells are translated into truncated proteins. We hypothesize that the acceleration of melanoma progression we observe in tumors expressing CDK13^mel^ could be influenced by the presence of these aberrant truncated proteins. Some of the proteins showing evidence of truncation in our data are tumor suppressors that would be expected to lose functionality; others may act in a dominant negative fashion to favor tumor progression^29–31^.

Recent reports suggest that prematurely terminated RNAs may be a newly recognized and more generalized oncogenic mechanism. Widespread intronic polyadenylation events were reported to disrupt tumor suppressor genes in leukemia by generating truncated mRNAs and proteins, however the truncation mechanism was not defined^38^. Loss of function mutations in CDK12 in metastatic castration-resistant prostate cancer and serous ovarian carcinomas were reported to cause increased production of truncated RNAs in DNA repair genes^28, 39^. The tumor suppressor function of CDK13 appears to uniquely function to target truncated messages for degradation via PAXT-dependent surveillance, representing an additional mechanism by which the oncogenic activity of prematurely terminated RNAs is countered in normal cells.

Our data are consistent with the evolutionarily related CDK12 and CDK13 having related but distinct functions (Table S1). Indeed, as known tumor suppressors their roles could be complementary, with CDK12 preventing the early termination of transcripts and CDK13 promoting the degradation of any prematurely terminated message that escape CDK12-mediated suppression.

CDK13 was recently identified as being required for proper development of neural crest-derived tissues. New truncating mutations in *CDK13* cause syndromic congenital heart disease^11^ and new kinase domain mutations identical to those we identified in melanoma cause a developmental disorder that affect heart, brain, and craniofacial development^10^. As melanoma is a cancer of neural crest derived melanocytes, neural crest derived tissues may have a particular sensitivity to CDK13 mutation, perhaps due to their broad differentiation potential and exquisite sensitivity to transcriptional changes. Our findings have prognostic implications for patients with CDK13-altered melanoma and suggest that patients with mutated or downregulated CDK13 should be considered for early stage adjuvant therapy trials. Our data suggest that patients with a CDK13-related developmental disorder should be monitored for melanoma.

## Methods

### Data Availability

The datasets generated during and/or analyzed during the current study are uploaded to GEO. The TCGA patient RNA-sequencing bams controlled-access data so they cannot be publically shared as they contain unique identifiers. The TCGA patient IDs used for analysis are listed in Supplementary Table 4.

### Code Availability

Custom code is available upon request.

### Patient mutation and survival analysis

Overall survival, mutation, and expression data (z-score values) from the TCGA melanoma cohort ^40^ were downloaded on 4/7/17 using cBioPortal^17^. A low-function CDK13 group was defined by either having a mutation in CDK13 or having low CDK13 expression, which was defined by z score equal to or less than −1.0. The difference in survival between low-function CDK13 vs. remaining cases was tested using a log-rank test. The difference in survival between low-function CDK13 stage 0, 1, or 2 patients were compared with remaining stage 1 or 2 patients. Mutation plot was downloaded on 4/7/17 and percent mutations were calculated with all data sets available 4/7/17 using cBioPortal^17^, excluding datasets from the Broad Institute, as it has duplicate samples with the DFCI/Broad set.

### Zebrafish model

All Experiments were performed in accordance with relevant guidelines and regulations. Animal studies were approved by Boston Children’s Hospital Institutional Animal Care and Use Committee (Protocol 17-10-3530R). Experiments were performed as published^19, 21^. Briefly, *p53^-/-^; mitfa:BRAFV600E;Na-/-* one-cell embryos were injected with either 20ng/uL control or experimental *MiniCoopR* (MCR) DNA along with tol2 *in vitro* transcribed RNA for integration. In overexpression experiments, control vectors expressed *EGFP*. In CRISPR experiments, control vectors expressed a gRNA to *arhgap11a*, a gene whose knock out has the same tumor curve as no gRNA vector in our system. In experiments where more than one vector was injected, DNA was prepared at 20ng/uL but divided equally between included vectors. For most experiments, embryos were sorted for melanocyte rescue at 5dpf. For *cdk13* CRISPR, control CRISPR, *CDK13 WT* expression, and *cdk13 CRISPR + CDK13 WT* expression embryos were sorted for melanocyte rescue, the 20 embryos with the most rescue were collected, and they were imaged at 3dpf. In all experiments, 20 zebrafish were raised per tank to control for density effects. Zebrafish were scored for the emergence of raised melanoma lesions as published^21^. Melanoma-free survival curves and Log-rank tests were generated in Prism. As melanocyte-specific *cdk13* CRISPR/Cas9 zebrafish and melanocyte-specific *CDK13 WT* expressing zebrafish had few melanocytes as adults, tumor curves were not gathered. All zebrafish regardless of sex were included as zebrafish sex is environmentally determined and not apparent until after most of the time points observed.

### Mutation Visualization on Crystal Structure

Patient mutations in *CDK13* were plotted on the crystalized kinase domain^18^ using PyMOL.

### *In vitro* Kinase Assays

Radioactive kinase reactions were performed using recombinant human CDK13 (694-1039) and Cyclin K (1-300) proteins. Purified recombinant wild type CDK13/CycK and mutant CDK13(R860Q)/CycK, CDK13(W878L)/CycK, and CDK13(K734R)/ CycK at a concentration of 0.2 µM were assayed for phosphorylation activity in a final volume of 30 µL containing kinase buffer (40 mM HEPES, pH 7.5, 34 mM NaCl, 34 mM KCl, 10 mM MgCl2, 5% glycerol), CDK13 substrates (GST-CTD f.l. 10 µM), cold ATP (to a final concentration of 1 mM) and 3 mCi [^32^P]-γ-ATP. The reaction mixture was incubated for 30 min at 30°C at 350 r.p.m. and terminated by adding EDTA to a final concentration of 50 mM. Aliquots of 15 µl each were spotted onto P81 Whatman paper squares. Paper squares were washed three times for 5 min with 0.75% (v/v) phosphoric acid, with at least 5 ml washing solution per paper square. Radioactivity was counted in a Beckman Scintillation Counter (Beckman-Coulter) for max. 5 min. Kinase assays were performed in duplicate and are represented as mean with s.d. One-way ANOVA for c-Myc substrate had an “F” value of 1084 and 4 degrees of freedom (dof). One-way ANOVA for CTD52 had an “F” value of 195 and 4 dof.

### Cloning (all except for mouse ES cells)

#### MiniCoopR (MCR) system^19, 21^

Zebrafish MCR overexpression constructs using the *mitfa* promoter, relevant coding sequence, 3’ pA tail, and the *MCR* destination vector were assembled with LR Clonase II Plus (Thermofisher, 12538120).

#### CDK13

Coding sequence was obtained using <u>HsCD00295573</u> from the Harvard Plasmid Repository (https://plasmid.med.harvard.edu/) and sequence verified. CDK13 *in vitro* mutagenesis for K734R, R860Q, P869S, W878L, P881L, and P893L were performed using primer design from www.genomics.agilent.com/primerDesignProgram.jsp, Q5 hot start polymerase with GC enhancer (NEB M0493S), and DpnI digest.

#### ZC3H14

Full-length nuclear isoform ZC3H14 coding sequence was obtained from Dharmacon (Accession: BC011793 Clone ID: 4298961). *In vitro* mutagenesis of S475D was done using the NEB site directed mutagenesis kit (NEB E0554S).

#### CRISPR/Cas9 Modified MCR system

The modified CRISPR MCR vector used as reported^19^. Briefly, CRISPR/Cas9 MCR constructs include a *mitfa* promoter driving Cas9 and a *U6* promoter driving expression of the gRNA. gRNAs to *cdk13, arhgap11a* (control), *ccnT1*, or *ccnK* were selected using CHOP CHOP^41, 42^ in the beginning of exons predicted to code for required protein domains. Prior to cloning into the *in vivo* system, multiple gRNAs were tested for cutting efficacy. To test cutting efficiency, *in vitro* synthesized gRNA [oligos were annealed and extended with T4 DNA polymerase and then synthesized with T7 RNA polymerase using the MEGAscript T7 kit (Thermofisher, AM1333)] and Cas9 protein (PNA Bio, CP01) were injected into one-cell stage zebrafish embryos. DNA was extracted at 48 hours using the HotSHOT method^43^, PCR across the gRNA site was done, and a T7E1 assay was used to assess cutting^41^. gRNAs with the most cutting by T7E1 were cloned into the MCR CRISPR vector using a BseRI and injected into the one cell stage. Subsequent workflows are described in other sections. CRISPR tumor analysis was done by extracting DNA from melanomas using the HotSHOT method^43^. PCR with a proofreading polymerase (NEB M0503S) was used to amplify across the cut site, and samples were submitted to the MGH DNA Core for CRISPR sequencing. Data sets were mapped to the *Danio rerio* genome (version GRCz11) using Bowtie (version 0.12.9). CrispR Variants Lite was used to identify insertions and deletions (indels) around the gRNA site (http://imlspenticton.uzh.ch:3838/CrispRVariantsLite/). Indels at the gRNA cut site in >1% of reads were used in downstream calculations. Reads that were predicted to maintain function included in-frame indels as well as wild type reads. Reads that were predicted to cause loss of function included out-of-frame indels.

### Zebrafish brightfield imaging

Zebrafish at 3 day, 2 week, 4 week, and 9 week timepoints were assessed for pigmentation and tumor onset via brightfield microscopy (Nikon DS-Ri2). Maximum backlight and LED illumination (NII-LED) settings were utilized with 0.5X magnification lens [1X/2X/3X zoom for 9/4/2 weeks post fertilization (wpf) zebrafish]. 3 day post fertilization (dpf) zebrafish were transferred to a 35mm dish, swirled, and photographed with about 20 fish in a photo at once with 0.5X magnification lens and 3X zoom. Melanocytes were quantified from 3dpf zebrafish photos and normalized to the number of zebrafish counted. One-way ANOVA was done. “F” = 13.14 and degrees of freedom = 3. Melanocyte patterns from 9wpf were categorized into a) no melanocytes, b) minimal melanocytes (0-33% of zebrafish length covered), c) strong melanocytes (34-100% of zebrafish length covered), or d) black patch (disrupts normal stripes and cross section is ≥ the distance between that zebrafishes eyes.). All zebrafish were anesthetized in tricaine solution (3-amino benzoic acidethylester, 4g/L) and oriented in an imaging mold (2% agarose in E3 buffer) prior to image capture.

### Human cell lines

A375 human melanoma cells were identity-verified (http://moleculardiagnosticscore.dana-farber.org/human-cell-line-identity-verification.html) and then used for transient transfections for IP-MS or knockdown of CDK13. Mycoplasma testing was done within one week of every experiment using human cell lines (Lonza, Mycoalert PLUS, LT07-710). All cell lines were always mycoplasma negative. Cells were grown in DMEM supplemented with 10% FBS, penicillin/streptomycin antibiotics, and L-glutamine.

### Expression for IP-mass spectrometry

LR Clonase II Plus was used to assemble pcDNA3.2 C-terminal V5 tag destination vector (ThermoFisher 12489019) with either *CDK13 WT, CDK13 W878L, CLOVER*, or *ZC3H14* in Constructs were transiently transfected using Lipofectamine 3000 per instructions (Thermofisher, L3000008).

### Stable Human Cell Lines

LR Clonase II Plus was used to assemble *CLOVER*, *CDK13 WT*, *CDK13 R860Q*, and *CDK13 W878L* into pLenti CMV Blast DEST. pLenti CMV Blast DEST (706-1)^44^ was a gift from Eric Campeau & Paul Kaufman (Addgene plasmid # 17451; http://n2t.net/addgene:17451; RRID:Addgene_17451). Lines were made in biologic triplicate. Cell counts were done at 24, 48, 72, and 96 hours. Cell doubling times calculated in exponential growth phase.

### Westerns in human melanoma cells

Protein was quantified using DC Protein Assay (Biorad, 5000116). For CDK13 westerns, 20μg was run on 3-8% tris acetate gel. Dry transfer was done at 25V for 10min. Primary anti-CDK13 antibody (Sigma HPA 059241, 1:1000) and primary loading control antibody anti-VCL (Sigma HPA 002131, 1:2500) were incubated overnight. For CLOVER, anti-GFP (Santa Cruz sc-9996, 1:1000) was incubated overnight. Rabbit or mouse secondary antibodies were incubated for 1hr at RT (CST 7074S or 7076S, 1:2000). Films were developed with Pierce ECL substrate (thermofisher, 32106).

### ChIP-sequencing

Zebrafish with melanomas expressing either *EGFP* or *CDK13 W878L* in *mitfa:BRAFmut+/+; p53-/-; Na-/-* zebrafish were sacrificed on ice and the tumors were dissected. 11 *EGFP* or 6 *CDK13 W878L* expressing melanomas were isolated at the same time point. Tumors were combined and minced. Cells were collected and fixed in 1.1% formaldehyde solution for 10min at room temperature (RT). Tumors were homogenized and filtered through a 100μm filter, spun down, washed with 1X PBS, and cells were counted using a Neubauer Chamber (9.00E+07 for *CDK13 W878L* and 7.10E+07 for *EGFP* expressing melanomas). Cells were spun again and the pellet was frozen with a dry ice/ethanol bath. ChIP was performed as published^45^, except upon lysis spike in *Drosphila* chromatin was added with 10ng per million cells (Active motif, 53083) and spike-in antibody 2μg was added to the beads (Active Motif, 61686). Briefly, IPs were performed using antibodies to CDK13 (Bethyl, A301-458A), hypophosphorylated RNAPII (abcam ab817, clone 8WG16, lot GR313984-17), RNAPII S2 CTD (ab5095; Lot G309257-1). Libraries were performed using the NEBNextMultiplex Oligos for Illumina kit (NEB) and run on an Illumina HiSeq 2000.

For ChIP-seq from human melanoma cells, one replicate (line 1) of A375 human melanoma cells expressing CLOVER and CDK13 R860Q as derived above in “**Stable Human Cell Lines**” were used. Equal cell number (approximately 40×10^6^) were used for each IP. Chromatin solubilization, spike in, and immunoprecipitation were performed using methods for zebrafish melanomas as above.

### ChIP-Seq processing and quantification

Raw reads were aligned twice: first to the dm6 revision of the D. melanogaster genome to remove exogenous spike-in reads using bowtie version 1.2.2 ^46^ with parameters -k 1 -m 1; unmapped reads were then aligned to the danRer10 revision of the zebrafish genome with default parameters or the non-random hg19 revision of the human genome with parameters -k 2 -m 2 and -l set to read length. Display files were created using MACS^47^ with parameters -w -S – space=50 –nomodel –shiftsize to get read counts in 50bp bins. These files were normalized to millions of mapped reads per sample and converted to TDF format using igvtools^48, 49^ toTDF.

Version 90 of the danRer10 Ensembl gene set was used for zebrafish ChIP-Seq analysis, and hg19 RefSeq gene positions were downloaded as part of the ROSE package (https://bitbucket.org/young_computation/rose/). Reads-per-million-normalized ChIP-Seq signal was quantified in promoters (transcription start sites +/- 250bp) using bamToGFF with parameters –m 1 –r –e −200 –f 1. Log2 fold-changes in signal were performed after adding one pseudocount to each condition. For displays of single-bin coverage, corresponding input signal calculated identically was subtracted from each region, and only regions with a positive input-subtracted CDK13 signal were shown.

Whole-gene metagenes across all genes greater than 2kb in length were constructed in three sections: −2kb upstream of the transcription start site to the transcription start site, transcription start site to the transcription end site, and transcription end site to 2kb downstream of the transcription end site. Matrices of coverage were calculated using bamToGFF (https://github.com/BradnerLab/pipeline) with parameters -m 50 -e 200 -r -f 1 for the upstream and downstream regions, and -m 150 -e 200 -r -f 1 for the genic region. The values in each bin were averaged across the genes used.

Gene lists were subsetted and/or broken into equally sized groups using different characteristics for different analyses. Unless otherwise noted, only genes greater than 2 kb from their transcription start sites to their transcription end sites were used.

Fold-change metagenes used the subset of all genes regardless of length that were in the 20% whose fold-changes in promoter CDK13 levels increased the most dramatically. The displayed metagenes represent per-bin log2-fold ratios of the average values for Ser2-RNA Polymerase II vs. the average values for total RNA Polymerase II where one pseudocount was added.

To study the effects of CDK13 mutation on coverage of genes expressed at different levels, the gene list was broken into six equally sized groups by their mean expression across three control samples for zebrafish samples. For human samples, the gene list was broken into five equally sized groups and expression amounts were determined from ribo-depleted RNA-sequencing from the three CLOVER expressing A375 lines as described above in “**Stable Human Cell Lines”.**

To study the effects of CDK13 mutation on coverage of genes of different lengths, genes were broken into six equally sized groups based on their gene length from transcription start site to transcription end site.

bigWig files displaying read coverage were created using readGAlignments and coverage from the GenomicAlignments^50^ package and export.bw from the rtracklayer ^51^ package, where putative PCR duplicate reads were removed. IGV (http://www.broadinstitute.org/igv) was used to view bigWig files in figures.

For anti-CDK13 ChIP-sequencing from melanomas from *mitfa:BRAFmut+/+; p53-/-; Na-/-* zebrafish expressing either *CDK13 R860Q + ccnT1* CRISPR or *CDK13 R860Q + ctrl* CRISPR, coverage of zebrafish promoters was quantified by bamToGFF. Gene positions for the GRCz10 revision of the zebrafish genome were downloaded from Ensembl (version 90). Two different windows were used: +/-2kb from the transcription start sites is displayed, and +/-250 bp from the transcription start sites was used to order rows/regions. Regions for display were broken into 100 equally sized bins (-m 100 -r), whereas ranking was done using a single window (-m 1 -r). In each case, input signal was subtracted from the corresponding regions. Promoters were subsetted for those that had background-subtracted RPM values in the 500 bp window > 1. They were ordered by their log2 fold-change between sgCtrl and sgccnT1 background-subtracted coverage, with one pseudocount added.

### Zebrafish Melanoma RNA-seq and RNA-seq processing

Zebrafish melanomas expressing EGFP, K734R, or R860Q in *mitfa:BRAFmut+/+; p53-/-; Na-/-* zebrafish were sacrificed on ice and the tumors were dissected, homogenized in RLT buffer, subjected to QIAshredder columns (Qiagen, 79656), and then RNA was isolated with a column based method with genomic DNA column removal (Qiagen, 74134). RNA was ribodepleted (Illumina Ribozero Gold, MRZG12324). Ribodepletion was confirmed with Agilent 4200 Tapestation. Library prepped with random priming (NEBNext Ultra RNA Library Prep Kit for Illumina, E7530), fragmentation time of 15min, 12 PCR cycles, and sequenced on an Illumina HiSeq 2000 (100bp paired end reads).

Sequenced reads were mapped to a custom version of the danRer10 transcriptome where all chromosomes were included plus human *CDK13* using tophat 2.1.1^52^ with parameters –library-type fr-unstranded –no-novel-juncs and -G set to a GTF of Ensembl zebrafish genes (v90). Expression quantification was performed using htseq-count^53^ using the same GTF as above and parameters -r name -i gene_name –stranded=no -f bam -m intersection-strict. Gene-level transcription read counts were then normalized to transcripts per million (TPM) (counts / bp in exons of all isoforms / 1000 / total counts across all genes / 1000000). Differential expression analysis was performed using DEseq2^54^ with standard parameters comparing raw read counts in all overexpression mutant samples against all overexpressed EGFP samples.

### Mouse Embryonic Stem Cells

#### Cell Culture, Cell Line Generation, Drug Conditions

All cell lines were tested for mycoplasma contamination periodically, including immediately after generation of CRISPR-modified clonal cell lines via the MycoAlert Mycoplasma Testing Kit (Lonza). Results were always negative for mycoplasma contamination. V6.5 (C57Bl/6-129) mESCs and derived cell lines were cultured on 0.2% gelatin-coated tissue culture plates in ES media: Dulbecco’s Modified Essential Media (Thermo Fisher) buffered with 10mM HEPES (Thermo Fisher) and supplemented with 15% Fetal Bovine Serum (Hyclone), 1000U/mL leukemia inhibitory factor (Millipore), 1x non-essential amino acids (Thermo Fisher), 2mM L-glutamine (Thermo Fisher), 0.11mM ß-mercaptoethanol (Sigma), 100 IU penicillin and 100ug/mL streptomycin (Corning).

Cdk13Δ clones were generated using CRISPR/Cas9 genome editing technology as follows (see Figure S3B). sgRNAs targeting intron 3 and intron 4 of the endogenous Cdk13 locus were cloned into pX458 (a gift from Feng Zhang, Addgene plasmid #48138)^55^ or pX330 (a gift from Feng Zhang, Addgene# 42230)^56^ respectively (see Supplementary Table. 3 for sequences). Two independent pairs of sgRNAs (one sgRNA targeting intron 3 and one sgRNA targeting intron 4) were co-transfected into wildtype V6.5 mouse embryonic stem cells with Lipofectamine®2000 (Thermo Fisher), and single-cell sorted for GFP+ fluorescence (transfected cells) 24 hours after transfection. Clones were screened for homozygous deletion of exon 4 by PCR and confirmed by sanger sequencing. Knockout of endogenous Cdk13 was confirmed by Western blot (Figures S3C). One knockout clone from each sgRNA pair was used throughout the study to control for off-target effects of the sgRNAs.

A Dox-inducible Cdk13 transgene was stably introduced into the two Cdk13Δ clones used throughout this studying using a piggybac retrotransposon system. N-terminal Flag-HA-tandem epitope-tagged Cdk13 (NP_001074527.1 isoform) was cloned via overlap extension PCR using two templates: (1) a synthetic gene block containing the codon-optimized N-terminus of Cdk13 (first 1577 base pairs) to reduce the high GC content in the region and (2) polyA-selected mouse cDNA from V6.5 cells. This PCR product was cloned into pCR8/GW/TOPO (Thermo Fisher) followed by transfer into the doxycycline-inducible piggybac expression vector, PBNeoTetO-Dest (a gift from A.W. Cheng), using standard TOPO and Gateway cloning kits (Thermo Fisher). The final Cdk13 transgene sequence is provided in Supplementary Table 5. This expression vector was cotransfected with pAC4 (constitutively expressing M2rtTA, the Dox-inducible transactivator, flanked by piggybac recombination sites, A.W. Cheng) and mPBase (piggybac transposase expression plasmid, A.W. Cheng) using Lipofectamine®2000. 24 hours after transfection, cells were selected with 150 ug/mL Hygromycin (Thermo Fisher) and 200 ug/mL G418 (Sigma) to select stable transformants and subsequently, single-cell cloned. Clones were screened for near wild-type levels of Cdk13 expression upon addition of 1ug/mL Dox for 24 hours. Two clones (one from each sgRNA pair used for knockout) were isolated that expressed near wild-type levels of Cdk13 expression upon addition of 1ug/mL Dox for 24 hours (representative Western blot for one clone shown in Figure S3E), and these two clones were used throughout the study.

#### Western Blotting

Whole cell extract was harvested by washing the cells in cold phosphate-buffered saline (PBS) and lysing in RIPA (10mM Tris pH7.4, 150mM NaCl, 1% TritonX-100, 0.1% SDS, 0.5% Sodium Deoxycholate, and 1mM EDTA) supplemented with 1x cOmplete, EDTA-free Protease Inhibitors (Roche) and 2uL/mL Benzonase Nuclease (Sigma). Lysates were incubated on ice for at least 30min, centrifuged for 10 min at 4°C at max speed, and the cleared lysate was quantified using a standard BCA assay (Thermo Fisher). Lysates were normalized for equivalent total protein in 1x Loading Dye (62.5mM Tris pH6.8, 5% glycerol, 2% SDS, 16.67% BME, and 0.083% bromophenol blue) or 1x NuPAGE LDS Sample Buffer (Thermo Fisher) with 1x NuPAGE Reducing Agent (Thermo Fisher). Normalized lysates were boiled for 5 min and run on NuPAGE 4-12% Bis-Tris Gels (Thermo Fisher). Gels were transferred overnight (30 V) to PVDF in 10% methanol supplemented 1x NuPAGE Transfer Buffer (NuPAGE Bis-Tris Gels). Primary antibodies used for blotting: Anti-HA High Affinity Antibody (Roche 11867423001), CDK13 (a gift from Arno L. Greenleaf), Enolase I (CST 3810S), alpha-Tubulin (Genescript a01410). Secondary antibodies used: ECL Anti-Rat IgG (GE Healthcare NA935V), ECL Anti-Mouse IgG (GE Healthcare NA931V), and ECL Anti-Rabbit IgG (GE Healthcare NA934V).

#### RNA Sequencing

Two independent Cdk13Δ clones with Dox-inducible Cdk13 were pre-treated with 1ug/mL Doxycycline (Dox) daily for at least 5 days prior to the start of the time course to express complementing levels of Cdk13 transgene in the Cdk13Δ background (Figure S3D). Dox was withdrawn from these cells at time 0, which resulted in significant knockdown and undetectable levels of Cdk13 after 48-or 72-hours respectively (Figure S3E). RNA was harvested in biological duplicate from both independent clones from cells maintained in Dox at time 0 (+Cdk13) or withdrawn from Dox for 48 or 72 hours using Trizol (Thermo). RNA was extracted following the standard Trizol protocol and subsequently DNase treated with Turbo DNase (Thermo Fisher) under standard reaction conditions. RNA quality was assessed by Agilent 2100 Bioanalyzer and only samples with a RIN value ≥ 8.9 were used for library preparation and sequencing. PolyA-selected libraries were made from 1ug of total RNA input using the TruSeq Stranded mRNA Library Prep Kit (Illumina RS-122-2102) with multiplexing barcodes, following the standard protocol with the following specifications: (1) 5 min RNA fragmentation time, (2) Superscript III (Thermo Fisher) was used for reverse transcription, (3) 15 cycles of PCR were used during the library amplification step, and (4) AMPure beads (Beckman Coulter) were used to size select/purify the library post PCR amplification instead of gel size selection. The 12 libraries were pooled and sequenced (75 base pair, paired-end reads) on one flow cell of an Illumina NextSeq500.

### 5’ to 3’ RNA-seq gradient analysis

For patient data, were approved via dbGaP for access to RNA-seq bams. RNA-seq bams were downloaded on 7/27/18 from cutaneous melanoma patients from The Cancer Genome Atlas (TCGA)^40^ from 3 patients with *CDK13* kinase-domain mutations and 5 best matched control *WT CDK13* patients by 1) stage, 2) age, 3) sex, and 4) oncogene from the National Cancer Institute (NCI) Genomic Data Commons (GDC) Data Portal (Patient characteristics in Supplementary Table 4). For zebrafish melanomas, bams from above for *CDK13 R860Q* (n=5) and EGFP (n=4) melanomas were used. For mouse ES cells, bams from above RNA-seq were used.

For each genome analyzed, The files were sorted using SAMtools^57^. All exons were assigned to one of the following classes: F: first exon, 5’-most exon in any transcript of the gene; AF: the first exon in an annotated transcript that is not the 5’-most exon in that gene; I: Internal exon, AL: the last exon in an annotated transcript that is not the 3’-most exon in that gene; L: last exon, 3’-most exon in all transcripts of the gene. The following annotations were used: human, Gencode version 28 gtf; zebrafish, GRCz10; mouse, custom annotation generated with custom scripts. For each transcript in the input annotation, the exons were sorted by position and the first exon in each transcript was assigned the value ‘AF’ and the last AL for “Alternative First/Last”. All others are I for “Internal”. After all transcripts belonging to a gene were processed, the complete list of exons was sorted by position to determine the 5’-most and 3’-most exons, which were assigned “F” and “L” status; all other exons retained their originally assigned classifications. All exons were then subjected to differential expression analysis using DEXSeq ^58, 59^. The log2-fold changes in exons belonging to each class were plotted and the differences in the distribution of these differential expression comparisons were determined using the non-parametric Wilcoxon rank sum test, two-sided.

IGV (http://www.broadinstitute.org/igv) was used to generate TDF files from human patient data and was used to view RNA in Figure 3E, signal normalized to 4^th^ exon as difference between internal exons did not change.

### Distal pA Transcripts per Million Analysis

Fastq files of reads from polyA-selected, strand-specific RNA sequencing that corresponded to the 3’ end of RNA fragments were first parsed to identify reads terminating in at least three adenosines. The putative polyA tails were then trimmed from the reads, and the remaining portion of the read was mapped, along with its mate pair, to the genome. Using custom scripts, properly mapped reads were then filtered by assessing whether all of the trimmed 3’ adenosines were genomically encoded, in which case they were removed from further analysis, or alternatively, represented evidence of posttranscriptional, non-templated polyadenylation in which case they were kept. Kept reads were then intersected using bedtools with a set of previously defined polyadenylation site clusters based on 3’ end sequencing from mouse embryonic stem cells of the same strain under steady-state conditions. Reads that overlapped known polyadenylation sites were then intersected with gencode annotated 3’ UTRs to identify polyA reads corresponding to the distal polyA site (full-length mRNA). Using Rsem/EBseq, the number of transcripts pertaining to each gene, normalized per million total transcripts estimated for each sample, was used as an estimate of relative transcript abundance. We calculated the ratio of read counts assigned as described above to the distal, full-length mRNA polyadenylation sites to the estimated transcripts per million (TPM) for each sample. The ratio of polyA reads/ transcript/ million for the four replicates of each genotype at each timepoint combined were then plotted as a box-and-whisker plot and the significance level of differences in population distributions were calculated by the Wilcoxon rank sum test.

### Tandem mass spectrometry

Protein was isolated from three separate EGFP and CDK13W878L zebrafish melanomas at week 25.9 weeks post fertilization in RIPA buffer with BME and they were submitted to the Thermo Fisher Scientific Center for Multiplexed Proteomics (TCMP) at Harvard (https://tcmp.hms.harvard.edu/) for tandem mass spectrometry with isobaric tags and synchronous precursor selection based MS3 technology. Peptides are selected for sequencing in MS1 scans. MS2 spectra are used for identifying peptides, and MS3 spectra are used for quantification via TMT reporter ions. MS2 spectra were searched using the SEQUEST algorithm against a Uniprot composite database derived from the Danio Rerio proteome containing its reversed complement and known contaminants. Peptide spectral matches were filtered to a 1% false discovery rate (FDR) using the target-decoy strategy combined with linear discriminant analysis. Proteins were quantified only from peptides with a summed SN threshold of >=100 and MS2 isolation specificity of 0.7. Using these parameters, 46,853 peptides with an FDR of <1% were identified which mapped to 6190 proteins.

### Tandem mass spectrometry data analysis

Data were filtered for peptides with unique identification. Peptide location was determined by dividing the peptide start location by the protein length and log2 fold change between CDK13W878L and EGFP was calculated. A D’Agostino and Pearson normality test (p=0.3374) showed data were normal. To plot proteome-wide changes, individual peptides that had a two-sided t-test p<0.1 were binned and plotted by % protein length (included 3676 measurements). Linear regression showed a slope of −0.5349 and the p=0.0001 that the slope was non zero.

To identify individual candidate proteins two methods were used. First, of the 3676 peptides identified above, proteins that had >1 measurement and a log2 fold changed slope of >-1 were included as candidates which identified 263 candidates. This method identified Separately, using all unique peptides, any protein with >3 peptide measurements were plotted for slope. Proteins with a negative slope were tested with an F test (p<0.05) to determine good fit. This method identified 103 candidates.

### IP-mass spectrometry

pcDNA3.2 plasmids expressing CLOVER, CDK13 WT, CDK13 W878L, or ZC3H14 were transfected using lipofectamine 3000 (Thermofisher L3000008) per protocol in 15cm2 plate in duplicate or triplicate into either A375s or *CDK13 R860Q* expressing A375s.. 48 hours after transfection, nuclei were isolated (Thermofisher 78833) and lysed per protocol with protease inhibitors with final volume of 500uL. 500uL of non-denaturing lysis buffer (NDLB = 20mM Tris pH8, 140mM NaCl, 10% glycerol, 1%triton, 5mM EDTA) with protease inhibitors (Sigma P8340) and phosphatase inhibitors [Na pyrophosphate (4mg/10mL) and Na fluoride (8mg/10mL)] was added. V5-conjugated agarose beads (Sigma A7345) were washed with phosphate buffered saline (PBS) per protocol and incubated overnight at 4C with lysate. Beads were washed twice with NDLB with protease inhibitors and phosphatase inhibitors and then four times with wash buffer with protease and phosphatase inhibitors (150mM NaCl, 50mM Tris pH 8). Protein was boiled in Laemmli Buffer, run on an 8% tris/glycine gel, stained with colloidal blue staining kit (Thermofisher LC6025), each lane was cut into 4 pieces, and each gel fragment was submitted for mass spectrometry using the Taplin Mass Spectrometry Facility at Harvard University. Proteins from CDK13 experiment were included in analysis if they were identified in all IP replicates. IP’ed proteins were excluded from analysis if they had >1 peptide in either control IP. Peptides for identified proteins were normalized as a percent of identified CDK13 peptides and only proteins that averaged at least 1% of CDK13 were plotted. *q values* were calculated with unpaired t test with multiple test correction using Prism software. Proteins from ZC3H14 experiment were included for analysis if they had at least 4 peptides in all replicates of one condition and >3x enrichment over control IP. We report here the 3 most significantly changed proteins and 2 unchanged proteins from the same complex. Statistics were done using multiple t-tests assuming similar scatter using Prism software.

ZC3H14 phosphorylation sites were reported if they were identified in all three biologic replicates with either 1) modification score >10 or 2) >2 peptides calling the same site.

### Zebrafish melanoma IHC

Zebrafish with melanomas were sacrificed on ice, fixed in 4%PFA 1xPBS 4-5d, washed in 1xPBS 2 days, washed in 50% ethanol 1 day, then stored in 70% ethanol. *CDK13 W878L* and *CDK13 P893L* melanomas were isolated at 18weeks post fertilization (wpf). *EGFP* melanomas were isolated at 18 and 19 weeks post fertilization because there were not enough tumors to isolate all of them at 18wpf. The first 10 melanomas with a gRNA to *arhgap11a* (control gRNA) were collected in batches as they arose (6 at 13.7wks and 4 at 20.3 weeks) or gRNA to *cdk13* (first three at 19weeks and the last three at 26.3weeks). Zebrafish were paraffin embedded, sectioned, and stained for PH3 Serine 10 (CST 9701) at 1:200 and then stained with secondary HRP-goat anti rabbit secondary (Dako K5007) by ServiceBio (http://www.servicebio.com). The square millimeter with the most PH3 positive cells was identified and then the PH3 positive cells were counted per square mm.

**Table 1:** CDK13 has a related but distinct biologic role and mechanism from CDK12.

|  | **CDK13** | **CDK12** |
| --- | --- | --- |
| **Mutated in Cancer** | Melanoma | Prostate <sup>60</sup>,<br>Ovarian <sup>61</sup> |
| **Mutated in Developmental Disorders** | New kinase domain mutations cause complex disorder involving several neural crest tissues<sup>10,62</sup>,<br>New truncation mutations cause syndromic congenital heart disease<sup>11</sup> | Not detected |
| **Target genes** | TNFalpha signaling | DNA repair <sup>63</sup><br>28,39 |
| **RNA Polymerase II** | Not required for RNAPII CTD S2 phosphorylation in melanoma | Required for RNAPII CTD S2 phosphorylation *in vivo* <sup>28</sup> |
| **RNA Mechanism** | Required for PAXT activation on prematurely terminated RNAs | Required to suppress intronic polyadenylation<br>28,39 |

## Acknowledgements

This research was supported by Damon Runyon Cancer Foundation Fellowship Award, American Society of Clinical Oncology Young Investigator Award, Charles S Memorial Sloan Kettering Starr Foundation Cancer Consortium, The Hope Funds for Cancer Research Grillo-Marxuach Family Fellowship, The American Lebanese Syrian Associated Charities (ALSAC), R01CA103846-14, and T32HL116324 from the National Heart, Lung, and Blood Institute. M.G. is a member of the excellence cluster ImmunoSensation (EXC2151) and funded by a grant from the Deutsche Forschungsgemeinschaft (GE 976/9-2). The CDK13 antibody used for mouse experiments was a gift from A.L. Greenleaf. The pAC4 and PBNeoTetO-Dest vectors were gifts from A. Cheng. S.D. was supported by a David H. Koch Fellowship.

## Author contributions

M.L.I. planned and performed the experiments with assistance from K.Y.C., T.F., and C.W. with the following exceptions. Mouse ES cell experiments were planned/performed by S.J.D. and kinase assays were planned/performed by S.D. M.L.I., B.J.A., P.B., S.B., and C.G.L. analyzed the data. D.L. coded the survival analysis. M.G. provided the plots of the mutations on the CDK13 crystal structure and insight into the effects the mutations could have on CDK13 function. K.A and T.H. advised on RNA experiments. R.A., P.S., P.B., and L.I.Z. supervised the project. M.L.I, B.J.A., P.B., and L.I.Z. wrote the manuscript.

## Materials and Correspondence

Correspondence and requests for materials should be addressed to L.I.Z.

## Extended Data

**Figure S1:**
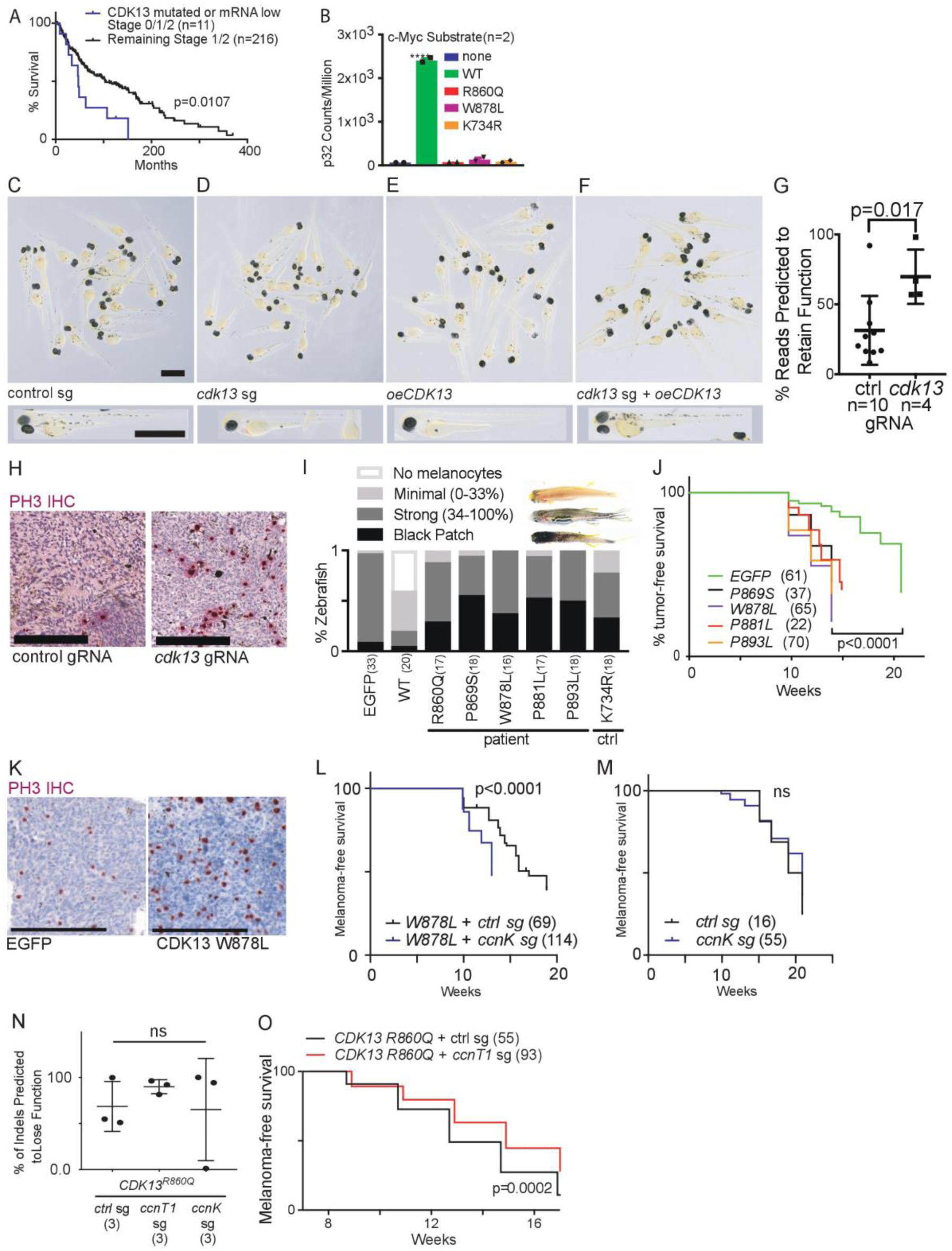
A) Downregulation or mutation of CDK13 in patients initially staged as 0/1/2 is associated with worse overall survival compared to remaining patients initially staged 1/2. p=0.0107. Log-rank. n= patients. B) *In vitro* kinase assay using wild type and patient-mutated CDK13 activated by CCNK on c-Myc substrate. One-way ANOVA with no kinase vs. all conditions; WT CDK13 ****=q=0.0001, all mutated CDK13 comparisons non-significant. Mean +/-SD. n=2 replicates. C-F) Light micrographs from 3 days post fertilization (dpf) *mitfa:BRAF*; p53-/-; *mitfa-/-* zebrafish injected with C) control melanocyte specific CRISPR, D) *cdk13* melanocyte-specific CRISPR, E) melanocyte-specific overexpression of human *CDK13^WT^*, F) *cdk13* melanocyte-specific CRISPR and human *CDK13^WT^*. Full image and inset scale bar = 100um. G) Representative PH3 staining image from *cdk13* gRNA and control gRNA zebrafish melanomas. Scale bar=200μm. H) Quantification of PCR reads across the CRISPR site predicted to retain function of *cdk13*. p=0.017 (t-test, two tailed). Mean +/- SD. n=melanomas. I) Quantification of pigmentation patterns of 9 week old *mitfa:BRAF*; p53-/-; mitfa-/- zebrafish from Figure 1H injected with melanocyte-specific expression vectors for *EGFP, CDK13^WT^, CDK13* patient mutations, or control catalytically dead *CDK13^K734R^*. White=no melanocytes, light gray = minimal melanocytes (0-33% length), dark gray = strong (34-100% length), black = black patch. (#)=zebrafish. J) Melanoma-free survival curves of EGFP and patient-mutant CDK13 melanomas. P<0.0001 (log rank). (#)=zebrafish. K) Representative PH3 staining image from *EGFP* and *CDK13^W878L^* expressing zebrafish melanomas. Scale bar=200μm. L) Melanoma-free survival of melanocyte-specific expression of *CDK13 W878L* coinjected with melanocyte-specific CRISPR of either a control gRNA or *ccnK* gRNA. p<0.0001 (log-rank). (#)=zebrafish. M) Melanoma-free survival of melanocyte-specific expression of control gRNA or *ccnK* gRNA. ns=non significant (log-rank). (#)=zebrafish. N) % CRISPR insertion/deletions (indels) predicted to lose function in *CDK13 R860Q* + control gRNA, *CDK13 R860Q* + *ccnT1* gRNA, and *CDK13 R860Q* + *ccnK* gRNA melanomas. Unpaired t-test, two tailed, non-significant. Mean +/-SD. (#)=melanomas. O) Melanoma-free survival of *mitfa:BRAF; p53-/-; mitfa-/-* zebrafish with melanocyte specific *CDK13 R860Q* expression and melanocyte-specific CRISPR of either control gRNA or *ccnT1* gRNA. p=0.0002, log rank. (#)=zebrafish.

**Figures S2:**
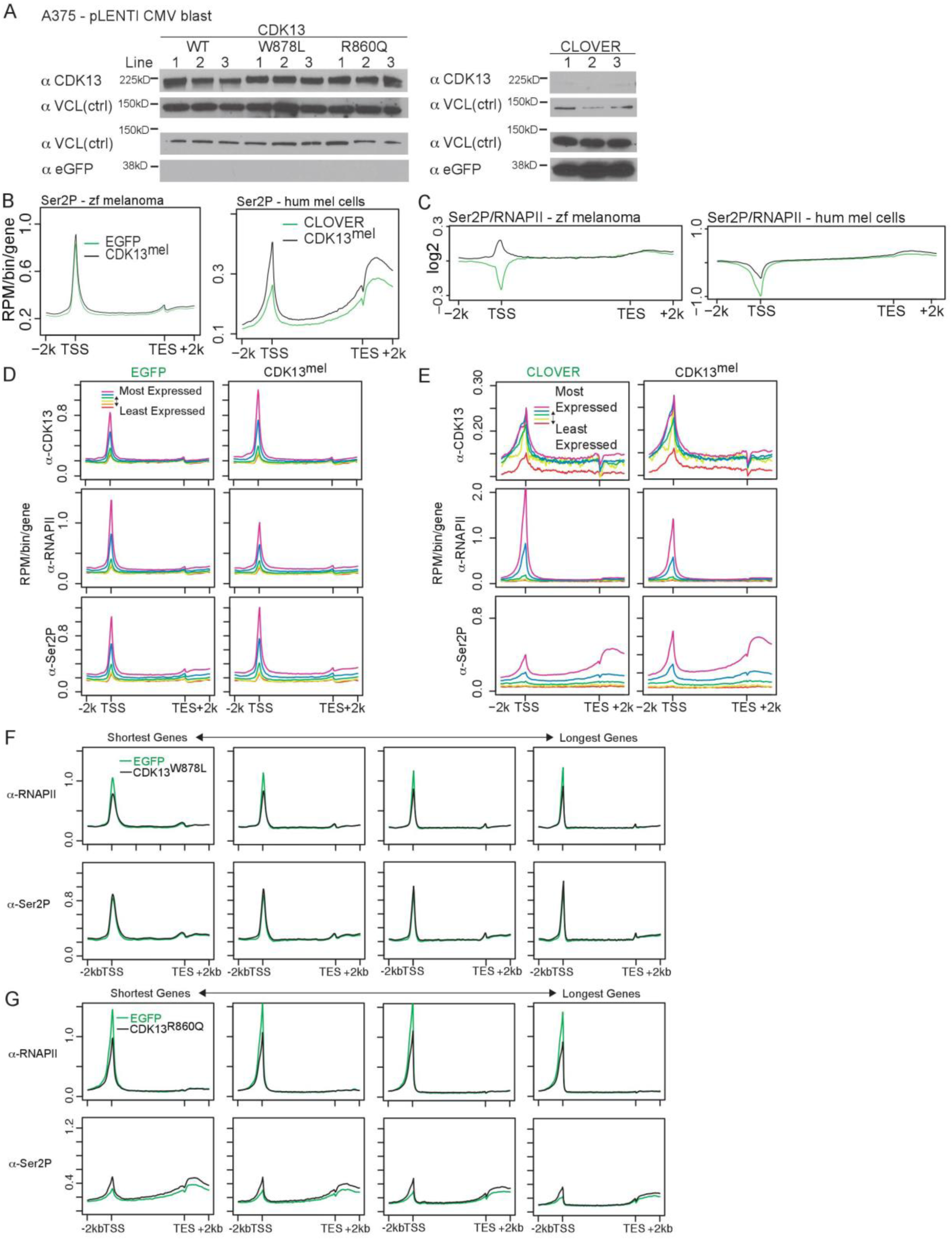
A) Immunoblot showing overexpression of CDK13 (WT, *W878L*, or *R860Q)* or CLOVER control in three independently derived lines. B) anti-RNAPII Ser2p metagene in zebrafish melanoma (left) and in human melanoma cells (right). C) ChIP-seq metagenes of Ser2p versus total RNAPII at genes with the largest fold increase in promoter CDK13^mel^ (left) in zebrafish melanoma and (right) in human melanoma cells. D-E) ChIP-seq metagenes for EGFP (left) and CDK13^mel^ (right) with anti-CDK13 (top), anti-8WG16 (middle), and anti-S2 CTD (bottom) in expression group sextiles. Purple = most highly expressed. Red = most lowly expressed for D) zebrafish melanomas and E) human melanoma cells. F-G) ChIP-seq metagenes for control (green) and mutant CDK13 (black) in gene length group sextiles with anti-RNAPII (top), and anti-S2 CTD (bottom) from F) zebrafish melanomas and in G) human melanoma cells. Longest and shortest gene groups removed for viewing ease. TSS = transcriptional start site. TES = transcriptional end site. 2k = 2kilobases. RPM = reads per million.

**Figure S3:**
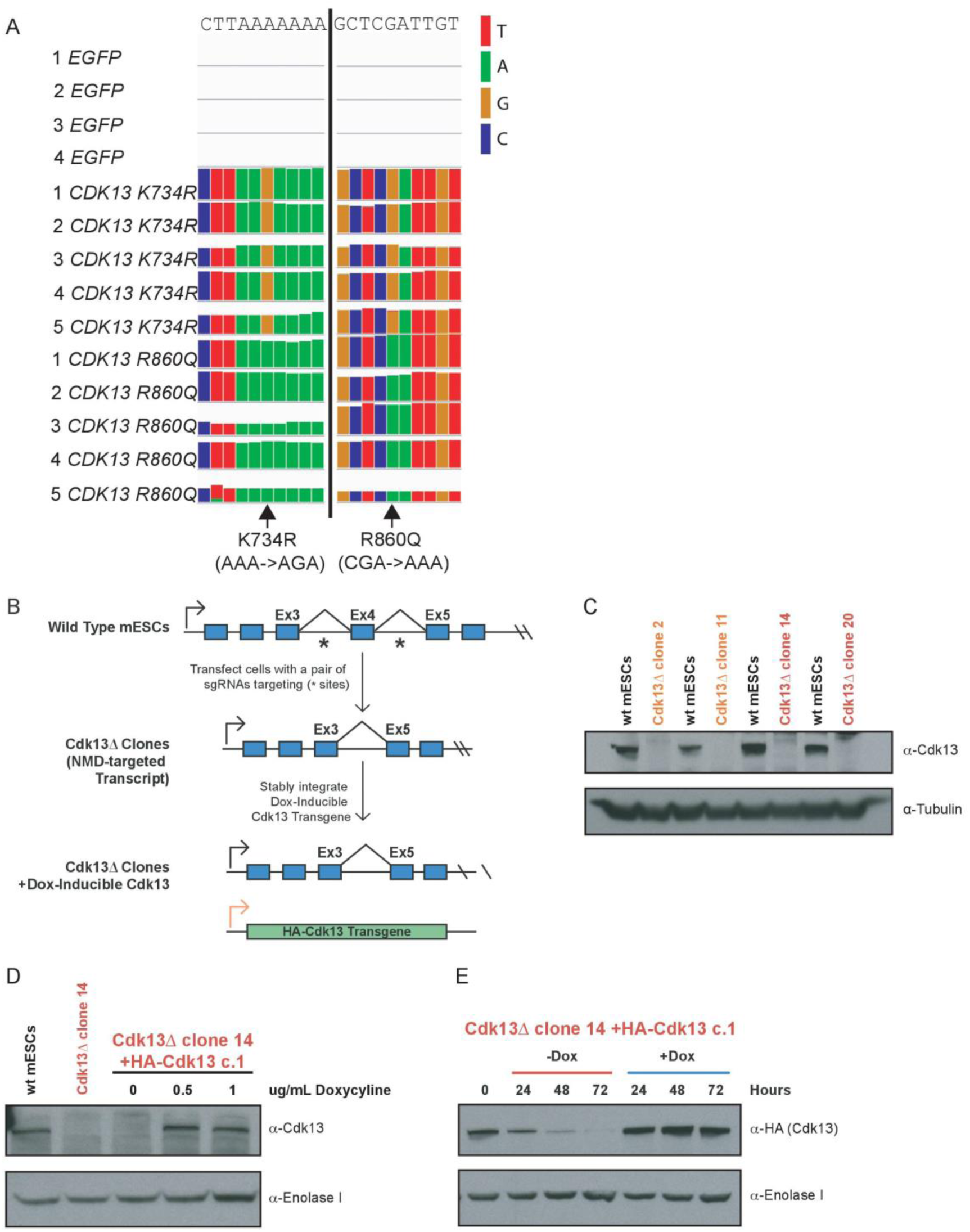
A) IGV track from zebrafish melanoma RNA-seq bam files confirming CDK13 mutant overexpression. Arrows=mutant base pairs. B) Schematic for generation of *Cdk13-/-* mouse embryonic stem cells (mESCs) with doxycycline-inducible *Cdk13-HA* for complementation. C) Representative immunoblot of n=4 showing 4 mESC clones with loss of *Cdk13*. Orange=gRNA pair 1. Red=gRNA pair 2. D) Representative immunoblot of n=2 (independently derived clones) showing near wild type level of rescue upon expression of *Cdk13-HA* upon expression of doxycycline. F) Representative immunoblot of n=2 (independently derived clones) of *Cdk13-HA* upon doxycycline withdrawal. RNA was gathered at 48 and 72hours.

**Figure S4:**
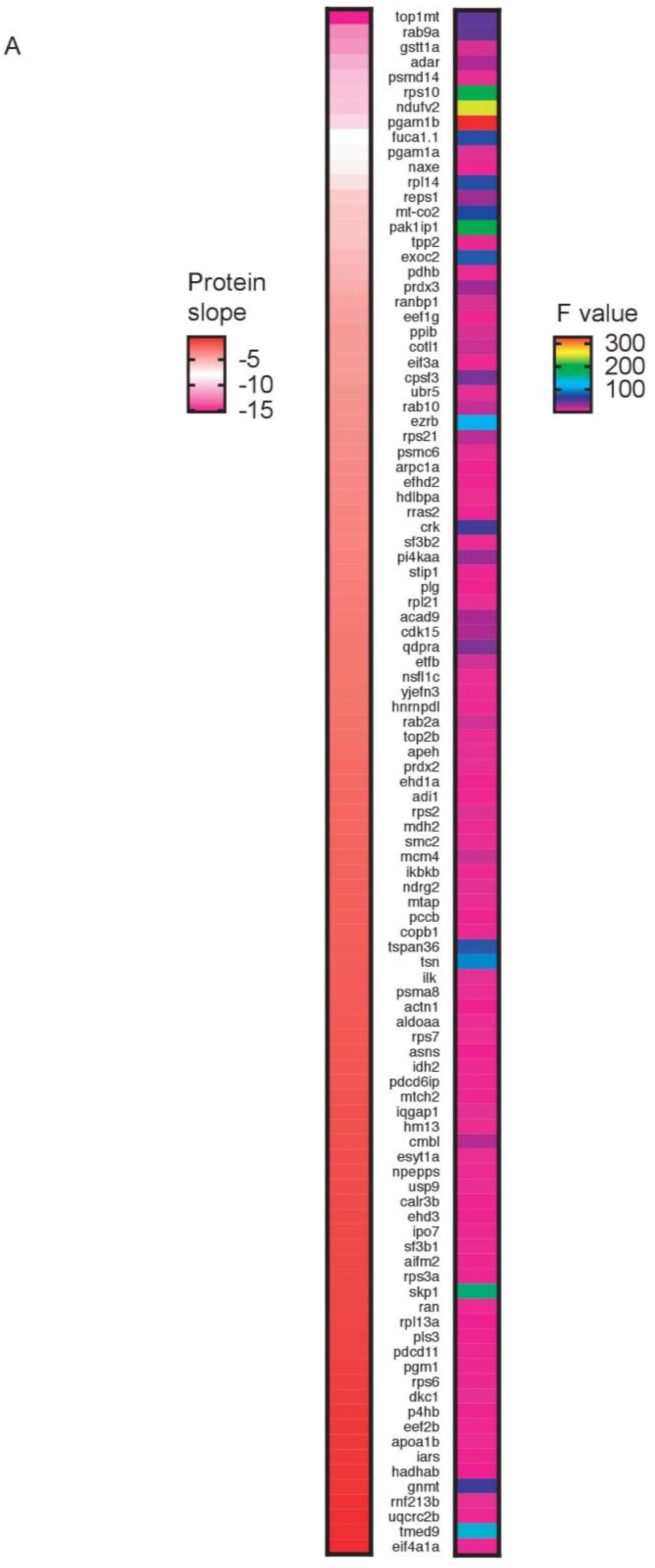
A) Heatmap of proteins with evidence of truncation in zebrafish CDK13^mel^ as compared with EGFP melanomas. Left = heatmap of log2 fold change slope measurements. Right = heatmap of F value (degree of significance, all included genes had F value indicating p<0.05).

**Figure S5:**
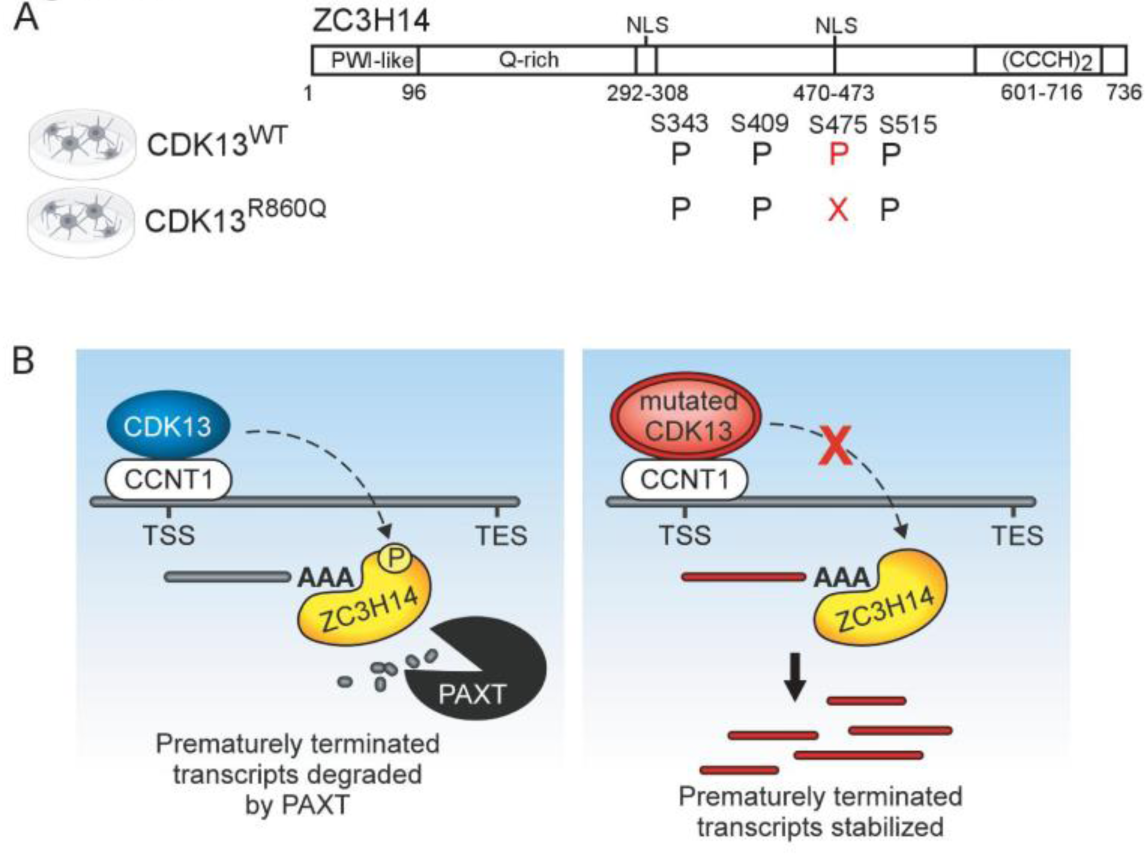
A) Graphic depicting ZC3H14 phosphorylations identified with in all three replicates of human melanoma cells with CDK13^WT^ or CDK13^R860Q^ expression. B) Model of CDK13’s activation of PAXT.

