## Supplemental Information File 1 for "*CDK13* Mutations Drive Melanoma via Accumulation of Prematurely Terminated Transcripts"

**Supplemental Information Table of Contents:**

1) DNA Oligos
2) Zebrafish gRNAs
3) Mouse gRNAs
4) TCGA Patient Characteristics
5) GEO Upload Files
6) Mouse Cdk13-HA rescue transgene

**Table 1: DNA Oligo Sequences**

| **Oligo Name** | **Sequence** |
|---|---|
| cdk13_gRNA5_assess_fw | catctttgtttcgttgttcagc |
| cdk13_gRNA5_assess_rev | tcaaaacacccaacactgaaag |
| cdk13_gRNA5_cloning_fw | ggagatcgtcaccgacaagggt |
| cdk13_gRNA5_cloning_rev | ccttgtcggtgacgatctccga |
| arhgap11a_gRNA_assess_fw | ttcaggcatttgttgctctg |
| arhgap11a_gRNA_assess_rev | ggagaagccgtgagtctgag |
| arhgap11a_gRNA_cloning_fw | ggagtcatgatgaccggaggt |
| arhgap11a_gRNA_cloning_rev | ctccggtcatcatgactccga |
| ccnT1_gRNA3_assess_fw | cgaatctgtgactgacaaggtc |
| ccnT1_gRNA3_assess_rev | acaacacgggacataaagaggt |
| ccnT1_gRNA3_cloning_fw | gagctgagtgtccaaagttagt |
| ccnT1_gRNA3_cloning_rev | taactttggacactcagctcga |
| ccnK_gRNA1_cloning_fw | ggtggtgaagcttccgaagggt |
| ccnK_gRNA1_cloning_rev | ccttcggaagcttcaccaccga |
| ccnK_gRNA1_assess_fw | agttcagactgatttgcggttt |
| ccnK_gRNA1_assess_rev | ggatacaatcccaaacgtgttc |
| CDK13_pENTR/D-TOPO_fw2 | caccatgccgagcagctcggaca |
| CDK13_pENTR/D-TOPO_revSTOP2 | tttagtatggtaaccctctgcctctgcctctgc |
| CDK13_pENTR/D-TOPO_revFUS2 | gtatggtaaccctctgcctctgcctctgc |
| CDK13_R860Q_fw | ttgcagactttggacttgctcaattgtatagctcagaagaaagtc |
| CDK13_R860Q_rev | gactttcttctgagctatacaattgagcaagtccaaagtctgcaa |
| CDK13_W878L_fw | tcaggtggacggtacaataaagtaattaccttgttagtatacg |
| CDK13_W878L_rev | cgtatactaacaaggtaattactttattgtaccgtccacctga |

| | |
|---|---|
| CDK13_P881L_fw | tactttatggtaccgtctacctgaactgctactgg |
| CDK13_P881L_rev | ccagtagcagttcaggtagacggtaccataaagta |
| CDK13_P869S_fw | gtaattaccttgttagtatacgaccgactttcttctgagctatac |
| CDK13_P869S_rev | gtatagctcagaagaaagtcggtcgtatactaacaaggtaattac |
| CDK13_P893L_fw | ctccatacatcaatggctagtgtgtatcgttcttctc |
| CDK13_P893L_rev | gagaagaacgatacacactagccattgatgtatggag |
| CDK13_R860Q_fw2_tm78 | ttcttctgagctatacaattgagcaagtccaaagtctgc |
| CDK13_R860Q_rev2_tm78 | gcagactttggacttgctcaattgtatagctcagaagaa |
| CDK13_K734R_for kinase assay | ccggcgagatggtggctctga**g**gaaagtgcgcctggacaacg |
| CDK13_R860Q_for kinase assay | ctggctgacttcggcctggctc**aa**ctgtactcctccgaggaa |
| CDK13_W878L_for kinase assay | ccaacaaagtgatcaccctgt**t**gtacaggccccctgagctgc |
| ZC3H14_pENTR/D-TOPO_fw | caccatggagatcggcaccgagatcag |
| ZC3H14_pENTR/D-TOPO_revFUS | ttcgctggtttgaggtcgaatcc |
| ZC3H14_S475D_fw | gccaaagctggatgaggaagtag |
| ZC3H14_S475D_rev | ttcttcagaataaatgatcttg |

**Table 2: Zebrafish CRISPR gRNAS (including cloning overhangs and PAMs)**

| **Gene Targeted** | **gRNA Sequence** |
|---|---|
| *arhgap11a* | GGGAGTCATGATGACCGGAGTGG |
| *cdk13* | GGAGATCGTCACCGACAAGGAGG |
| *ccnT1* | GAGCTGAGTGTCCAAAGTTAAGG |
| *ccnK* | GGTGGTGAAGCTTCCGAAGG |

**Table 3: Mouse CRISPR gRNAS (not including overhangs or PAMs)**

| **Gene Targeted** | **Location** | **gRNA Sequence** |
|---|---|---|
| *Cdk13* (pair 1) | Intron 3 | GTATCTTCCTGCTAAATATC |
| | Intron 4 | AAGGGTAAACTTCTATACAT |
| *Cdk13* (pair 2) | Intron 3 | AAGACACTAATCTATCATTA |
| | Intron 4 | TGTATCACCAGCCCTCAAGA |

**Table 4: TCGA Melanoma Patient Characteristics**

| ID | CDK13 Status | Sex | Age | Initial Stage | Oncogene |
|---|---|---|---|---|---|
| TCGA-FS-A4FD | R860Q | M | 39 | T2N3, Stage IIIC | NRAS |
| TCGA-EE-A2MD | P881L | M | 52 | T3aN0, Stage IIA | NRAS |
| TCGA-EE-A29C | I843N | M | 20 | T2a, Stage Ib | BRAF |
| TCGA-EE-A2GS | WT | F | 28 | T2a, Stage Ib | BRAF |
| TCGA-D3-A3MV | WT | F | 38 | T2bN2a, Stage IIIb | NRAS |
| TCGA-FS-A1ZP | WT | M | 52 | T3N0, Stage II | NRAS |
| TCGA-EE-A2MG | WT | M | 23 | T2N0, Stage I | BRAF |
| TCGA-DA-A3F8 | WT | M | 39 | T2aN2b, Stage IIIb | BRAF |

### Table 5: GEO Upload Files

| File Name | File Type | Assay | Organism |
|---|---|---|---|
| A375_CMV_PE_CDK13_2.fastq.gz<br>A375_CMV_PE_input_2.fastq.gz<br>A375_CMV_PE_CDK13_1.fastq.gz<br>A375_CMV_PE_input_1.fastq.gz<br>A375_CMVCDK13_SE_Ser2PolII_2.fastq.gz<br>A375_CMVCDK13_SE_Ser2PolII_1.fastq.gz<br>A375_CMVCDK13_PE_input_1.fastq.gz<br>A375_CMVCDK13_PE_input_2.fastq.gz<br>A375_CMVCDK13_PE_CDK13_1.fastq.gz<br>A375_CMVCDK13_PE_CDK13_2.fastq.gz<br>A375_CMV_SE_input_2.fastq.gz<br>A375_CMVCDK13_SE_PolII_2.fastq.gz<br>A375_CMVCDK13_SE_PolII_1.fastq.gz<br>A375_CMV_SE_input_1.fastq.gz<br>A375_CMV_SE_Ser2PolII_1.fastq.gz<br>A375_CMV_SE_Ser2PolII_2.fastq.gz<br>A375_CMV_SE_PolII_1.fastq.gz<br>A375_CMV_SE_PolII_2.fastq.gz<br>A375_CMVCDK13_SE_input_1.fastq.gz<br>A375_CMVCDK13_SE_input_2.fastq.gz | Raw | ChIP-seq | *Homo sapiens* |
| A375_CMV_PE_CDK13_comb.bw<br>A375_CMV_PE_input_comb.bw<br>A375_CMV_SE_input_comb.bw<br>A375_CMV_SE_PolII_comb.bw<br>A375_CMV_SE_Ser2PolII_comb.bw<br>A375_CMVCDK13_PE_CDK13_comb.bw<br>A375_CMVCDK13_PE_input_comb.bw<br>A375_CMVCDK13_SE_input_comb.bw<br>A375_CMVCDK13_SE_PolII_comb.bw | Processed | ChIP-seq | *Homo sapiens* |

| | | | |
|---|---|---|---|
| A375_CMVCDK13_SE_Ser2PolII_comb.bw | | | |
| MI_RNA_1_USPD16093847-5_HW57WBBXX_L7_1.fq.gz<br>MI_RNA_1_USPD16093847-5_HW57WBBXX_L7_2.fq.gz<br>MI_RNA_2_USPD16093847-6_HW57WBBXX_L7_1.fq.gz<br>MI_RNA_2_USPD16093847-6_HW57WBBXX_L7_2.fq.gz<br>MI_RNA_3_USPD16093847-7_HW57WBBXX_L7_1.fq.gz<br>MI_RNA_3_USPD16093847-7_HW57WBBXX_L7_2.fq.gz | Raw | RNA-seq | *Homo sapiens* |
| RNA1.tophat.htseq-count.out<br>RNA2.tophat.htseq-count.out<br>RNA3.tophat.htseq-count.out | Processed | RNA-seq | *Homo sapiens* |
| CDK13_R860Q_oe_R161.L001_R1.fastq.gz<br>CDK13_R860Q_oe_R161.L001_R2.fastq.gz<br>CDK13_R860Q_oe_R161.L002_R1.fastq.gz<br>CDK13_R860Q_oe_R161.L002_R2.fastq.gz<br>CDK13_R860Q_oe_R160.L001_R1.fastq.gz<br>CDK13_R860Q_oe_R160.L001_R2.fastq.gz<br>CDK13_R860Q_oe_R160.L002_R1.fastq.gz<br>CDK13_R860Q_oe_R160.L002_R2.fastq.gz<br>EGFP_oe_R165.L001_R1.fastq.gz<br>EGFP_oe_R165.L001_R2.fastq.gz<br>EGFP_oe_R165.L002_R1.fastq.gz<br>EGFP_oe_R165.L002_R2.fastq.gz<br>EGFP_oe_R164.L001_R1.fastq.gz<br>EGFP_oe_R164.L001_R2.fastq.gz<br>EGFP_oe_R164.L002_R1.fastq.gz<br>EGFP_oe_R164.L002_R2.fastq.gz<br>EGFP_oe_R163.L001_R1.fastq.gz<br>EGFP_oe_R163.L001_R2.fastq.gz<br>EGFP_oe_R163.L002_R1.fastq.gz<br>EGFP_oe_R163.L002_R2.fastq.gz<br>EGFP_oe_R162.L001_R1.fastq.gz<br>EGFP_oe_R162.L001_R2.fastq.gz<br>EGFP_oe_R162.L002_R1.fastq.gz<br>EGFP_oe_R162.L002_R2.fastq.gz<br>CDK13_R860Q_oe_R157.L001_R1.fastq.gz<br>CDK13_R860Q_oe_R157.L001_R2.fastq.gz<br>CDK13_R860Q_oe_R157.L002_R1.fastq.gz<br>CDK13_R860Q_oe_R157.L002_R2.fastq.gz<br>CDK13_K734R_oe_R152.L001_R1.fastq.gz<br>CDK13_K734R_oe_R152.L001_R2.fastq.gz<br>CDK13_K734R_oe_R152.L002_R1.fastq.gz<br>CDK13_K734R_oe_R152.L002_R2.fastq.gz<br>CDK13_K734R_oe_R153.L001_R1.fastq.gz<br>CDK13_K734R_oe_R153.L001_R2.fastq.gz<br>CDK13_K734R_oe_R153.L002_R1.fastq.gz<br>CDK13_K734R_oe_R153.L002_R2.fastq.gz<br>CDK13_K734R_oe_R154.L001_R1.fastq.gz<br>CDK13_K734R_oe_R154.L001_R2.fastq.gz<br>CDK13_K734R_oe_R154.L002_R1.fastq.gz<br>CDK13_K734R_oe_R154.L002_R2.fastq.gz<br>CDK13_K734R_oe_R155.L001_R1.fastq.gz<br>CDK13_K734R_oe_R155.L001_R2.fastq.gz<br>CDK13_K734R_oe_R155.L002_R1.fastq.gz<br>CDK13_K734R_oe_R155.L002_R2.fastq.gz<br>CDK13_R860Q_oe_R158.L001_R1.fastq.gz<br>CDK13_R860Q_oe_R158.L001_R2.fastq.gz | Raw | RNA-seq | *Danio rerio* |

| | | | |
|---|---|---|---|
| CDK13_R860Q_oe_R158.L002_R1.fastq.gz<br>CDK13_R860Q_oe_R158.L002_R2.fastq.gz<br>CDK13_K734R_oe_R156.L001_R1.fastq.gz<br>CDK13_K734R_oe_R156.L001_R2.fastq.gz<br>CDK13_K734R_oe_R156.L002_R1.fastq.gz<br>CDK13_K734R_oe_R156.L002_R2.fastq.gz<br>CDK13_R860Q_oe_R159.L001_R1.fastq.gz<br>CDK13_R860Q_oe_R159.L001_R2.fastq.gz<br>CDK13_R860Q_oe_R159.L002_R1.fastq.gz<br>CDK13_R860Q_oe_R159.L002_R2.fastq.gz | | | |
| CDK13_R860Q_oe_R161.htseq.txt<br>CDK13_R860Q_oe_R160.htseq.txt<br>EGFP_oe_R165.htseq.txt<br>EGFP_oe_R164.htseq.txt<br>EGFP_oe_R163.htseq.txt<br>EGFP_oe_R162.htseq.txt<br>CDK13_R860Q_oe_R157.htseq.txt<br>CDK13_K734R_oe_R152.htseq.txt<br>CDK13_K734R_oe_R153.htseq.txt<br>CDK13_K734R_oe_R154.htseq.txt<br>CDK13_K734R_oe_R155.htseq.txt<br>CDK13_R860Q_oe_R158.htseq.txt<br>CDK13_K734R_oe_R156.htseq.txt<br>CDK13_R860Q_oe_R159.htseq.txt | Processed | RNA-seq | *Danio rerio* |
| EGFP_Ser2PolII.fastq.gz<br>EGFP_PolII.fastq.gz<br>EGFP_input.fastq.gz<br>EGFP_CDK13.fastq.gz<br>CDK13_W878L_Ser2PolII.fastq.gz<br>CDK13_W878L_PolII.fastq.gz<br>CDK13_W878L_input.fastq.gz<br>CDK13_W878L_CDK13.fastq.gz | Raw | ChIP-seq | *Danio rerio* |
| EGFP_Ser2PolII.bw<br>EGFP_PolII.bw<br>EGFP_input.bw<br>EGFP_CDK13.bw<br>CDK13_W878L_Ser2PolII.bw<br>CDK13_W878L_PolII.bw<br>CDK13_W878L_input.bw<br>CDK13_W878L_CDK13.bw | Processed | ChIP-seq | *Danio rerio* |
| 150927_C5FHE_1572A_L1_1_Library19_sequence.fastq<br>150927_C5FHE_1572A_L1_2_Library19_sequence.fastq<br>150927_C5FHE_1572A_L1_1_Library20_sequence.fastq<br>150927_C5FHE_1572A_L1_2_Library20_sequence.fastq<br>150927_C5FHE_1572A_L1_1_Library21_sequence.fastq<br>150927_C5FHE_1572A_L1_2_Library21_sequence.fastq<br>150927_C5FHE_1572A_L1_1_Library22_sequence.fastq<br>150927_C5FHE_1572A_L1_2_Library22_sequence.fastq<br>150927_C5FHE_1572A_L1_1_Library23_sequence.fastq<br>150927_C5FHE_1572A_L1_2_Library23_sequence.fastq<br>150927_C5FHE_1572A_L1_1_Library24_sequence.fastq | Raw | RNA-seq | *Mus musculus* |

| |
|---|
| 150927_C5FHE_1572A_L1_2_Library24_sequence.fastq<br>150927_C5FHE_1572A_L1_1_Library25_sequence.fastq<br>150927_C5FHE_1572A_L1_2_Library25_sequence.fastq<br>150927_C5FHE_1572A_L1_1_Library26_sequence.fastq<br>150927_C5FHE_1572A_L1_2_Library26_sequence.fastq<br>150927_C5FHE_1572A_L1_1_Library27_sequence.fastq<br>150927_C5FHE_1572A_L1_2_Library27_sequence.fastq<br>150927_C5FHE_1572A_L1_1_Library28_sequence.fastq<br>150927_C5FHE_1572A_L1_2_Library28_sequence.fastq<br>150927_C5FHE_1572A_L1_1_Library29_sequence.fastq<br>150927_C5FHE_1572A_L1_2_Library29_sequence.fastq<br>150927_C5FHE_1572A_L1_1_Library30_sequence.fastq<br>150927_C5FHE_1572A_L1_2_Library30_sequence.fastq |
