## Supplemental Information File 2 for "*CDK13* Mutations Drive Melanoma via Accumulation of Prematurely Terminated Transcripts"

ACTACTGGTACCGCCGCCACCATGGACTACAAGGACGACGATGACAAGTACCCTTATGACGTGCCCGAT
TACGCTCCTTCCAGCTCCGATACAGCTCTCGGCGGAGGAGGCGGACTCAGCTGGGCCGAGAAAAAGCT
GGAGGAGAGGAGGAAGAGGAGGAGGTTTCTCAGCCCTCAGCAGCCTCCTCTGCTGCTCCCTCTCCTGCA
ACCTCAGCTCCTCCAACCCCCTCCTCCTCCCCCTCCTCTCCTGTTTCTGGCTGCTCCTGGAGCCGCTGCCGC
TGCCGCCGCTGCTGCTGCCGCTTCCAGCTCCTGTTTTAGCCCTGGCCCTCCTCTGGAGGTCAAGAGGCTG
GCCAGAGGAAAAAGGAGACCCGGCGGCAGGCAAAAGAGAAGGAGAGGACCTAGGGCCGGACAGGAA
GCCGAAAAGAGGAGAGTGTTCTCCCTGCCTCAACCCCAGCAGGATGGAGGCGGAGGAGCTTCCAGCGG
AGGAGGAGTGACACCCCTGGTGGAGTACGAGGACGTGTCCAGCCAAAGCGAACAAGGCCTGCTCCTGG
GAGGAGCTAGCGCTGCTACAGCTGCCACCGCTGCTGGAGGAACCGGAGGAAATGGAGGATCCCCCGCT
TCCAGCAGCGGAACCCAGAGAAGAGCCGAGGGCAGCGAGAGAAGACCTAGAAGGGACAGAAGGAGC
TCCAGCGGCAGGTCCAAAGAGAGGCACAGGGAACATAGGAGGAGAGATGGCACAAGGAGCGGATCC
GAGGCCAGCAAGGCCAGATCCAGACACGGACACAGCGGCGAGGAAAGAGCCGAGGCTGCCAAATCCG
GCTCCTCCAGCTCCTCCGGCGGAAGAAGAAAGTCCGCCTCCGCCACATCCTCCAGCAGCAGCTCCAGGA
AGGACAGGGACCTCAAGGCTCATAGGAGCAGGACCAAGAGCTCCAAGGAACCCCCTTCCGCCTACAAG
GAGCCCCCCAAAGCCTACAGGGAAGACAAGTCCGAACCCAAGGCCTACAGGAGGAGGCAGAGGTCCCT
CAGCCCTCTCGGAGGCAGAGACGAGTCCCCTGTCAGCCACAGGGCTAGCCAGTCCCTGAGATCCAGGA
AAAGCCCTTCCCCCGCTGGAGGCGGAAGCAGCCCTTATTCCAGAAGACTGCCCAGGTCCCCTAGCCCCT
ACAGCAGGAGGAGATCCCCCAGCTACAGCAGGCATTCCTCCTACGAGAGAGGAGGAGATGTCTCCCCC
TCCCCCTACAGCTCCTCCTCCTGGAGGAGGAGCAGATCCCCCTATTCCCCTGTGCTGAGGAGGTCCGCCA
AGTCCAGGTCCAGGAGCCCTTACTCCTCCAGGCACAGCAGAAGCAGGTCCAGGCACAGGCTGAGCAGG
TCCAGAAGCAGGCACTCCTCCATCAGCCCCTCCACCCTGACACTGAAGAGCTCCCTGGCCGCCGAGCTCA
ACAAGAACAAGAAAGCTAGGGCCGCTGAGGCTGCTAGAGCTGCCGAAGCCGCTAAAGCCGCCGAAGCC
GCCAAAGCTGCTGAAGCCGCCGCCAAGGCTGCTAAGGCCTCCAACGCTTCCACACCCACCAAGGGAAAC
ACCGAGACCGGAGCCTCCGTCTCCCAGACCAACCACGTGAAGGAAGTCAAGAAGCTTAAAACTGAGCAT
GCACCTTCTCCTTCAAGTGGTGGGACCGTCAAAAGCGACAAAGCAAAAACAAAGCCACCGCTTCAAGTA
ACAAAGGTAGACAATAATTTGACAGTAGAGAAAGCCACCAAGAAAACAGTTGTTGGGAAGGAGAGTAA
ACCTGCTGCTACAAAGGAAGAACCAGTTTCCACTAAAGAGAAAAGCAAGCCACTCACACCAAGCACAGG
AGCCAAGGAGAAGGAGCAGCATGTGGCTTTAGTGACCTCTACGTTACCGCCATTACCTTTGCCTCCCATG
CTGCCTGAAGATAAAGATGCTGATAGCTTAAGAGGCAACATTTCTGTCAAAGCAGTTAAAAAAGAAGTA
GAAAAGAAACTCCGATGTCTGCTTGCTGATTTACCATTGCCCCCTGAGTTACCAGGAGGAGATGATCTTT
CCAAGAGTCCAGAGGAGAAGAAAACAGCAGCACAGTTACATAGCAAACGAAGGCCTAAAATATGTGGG
CCTCGCTATGGTGAAATCAAAGAAAAAGATATTGACTGGGGGAAACGCTGCGTGGATAAATTTGATATC
ATCGGAATTATTGGAGAAGGTACTTATGGACAAGTTTACAAAGCCAGGGACAAAGACACGGGAGAAAT
GGTAGCCTTAAAGAAAGTACGTCTGGATAATGAAAAGGAGGGTTTCCCAATTACAGCAATTAGAGAAAT
TAAAATTCTTCGGCAACTCACCCACCAGAGTATCATCAATATGAAGGAAATCGTGACTGATAAAGAAGA
TGCTTTGGATTTTAAGAAAGACAAAGGTGCATTTTACCTGGTGTTTGAATATATGGACCATGATCTGATG
GGACTGCTGGAATCAGGCTTGGTTCATTTTAATGAAAACCATATAAAATCTTTTATGAGACAGCTCATGG
AAGGCCTGGATTATTGTCATAAGAAGAACTTTTTGCATAGAGATATTAAATGTTCAAATATCCTTCTAAAT
AATAGAGGACAGATAAAACTTGCAGATTTTGGACTTGCTCGTTTATATAGCTCAGAAGAAAGTCGCCCA
TATACTAACAAGGTCATTACTTTGTGGTATCGTCCACCTGAATTGCTCTTGGGAGAAGAACGATATACAC
CAGCCATTGATGTATGGAGCTGTGGATGTATCCTTGGTGAACTCTTCACTAAAAAACCTATATTTCAAGC
AAACCAGGAACTTGCACAGCTAGAGCTAATAAGCCGTATATGTGGGAGTCCATGTCCTGCAGTGTGGCC
TGATGTAATCAAACTGCCATATTTCAACACCATGAAACCAAAGAAGCAATATCGGCGGAAGTTAAGAGA
AGAATTTGTTTTCATCCCCGCAGCTGCACTCGACTTATTCGATTACATGCTTGCCTTGGATCCCAGTAAGC

GCTGCACTGCTGAGCAGGCTCTTCAGTGTGAGTTCCTGCGAGACGTGGAACCCTCCAAAATGCCTCCAC
CAGACCTTCCTTTGTGGCAAGATTGTCATGAATTATGGAGTAAAAAGAGAAGAAGACAGAAACAGATG
GGCATGACTGATGATCTTTCCACAATCAAAGCCCCTAGGAAGGACCTGTCTCTGGGCTTAGATGACAGC
AGAACTAACACACCCCAGGGTGTGCTGCCACCCGCACAGTTGAAATCTCAGAGCAACTCAAATGTAGCA
CCTGTAATAACAGGCCCTGGACAGCCGTTAAACCACAGTGAATTGGCAATTCTTCTAAACCTACTACAAT
CTAAATCAAGTGTTAATATGGCTGATTTTGTCCAAGTGTTGAACATTAAGGTAAACTCTGAGACTCAACA
GCAGCTAAATAAAATAAACCTTCCTGCTGGAATTTTGGCAACAGGTGAAAAACAGACAGATCCATCAAC
ACCACAACAGGAGTCTTCAAAATCATTGGGAGGAGTTCAGCCTTCACAGACCATCCAGCCTAAAGTGGA
AACTGATGCTGCCCAGGCTGCTGTGCAGAGTGCATTTGCAGTCCTCTTGACTCAGTTAATAAAGGCCCAA
CAGTCCAAACAGAAAGATGCCATGCTAGAGGAAAGGGAAAATGGATCAGGACATGAAGCTCCATTGCA
ACTCAGGCCTCCTCTAGAACCGAGCACTCCTGGATCCGGGCAAGATGACCTCATCCAGCACCAAGACAG
GAGGATATTGGAGCTGACACCAGAACCAGACCGGCCTCGAATTCTGCCTCCTGATCAACGGCCTCCTGA
ACCTCCTGAACCACCACCAGTTACTGAGGAGGACCTAGATTATCGGACAGAAAACCAGCATGTTCCTACC
ACTAGTTCTTCATTAACTGACCCACATGCTGGAGTGAAGGCAGCCCTCTTACAGCTGCTTGCTCAGCATC
AGCCCCAGGATGATCCCAAAAGAGAAGGTGGTATCGATTATCCCACAGGAGACACATATGTGCCCAGTT
CAGACTATAAGGACAACTTTGGATCTTCTTTCTCTGCCGCTCCTTACGTTAGCAGTGATGGTCTAGGAAG
TAGCTCCGCTGCTGCACCATTGGAAGCACGTAGTTTCATTGGAAACTCAGATATTCAGTCTCTGGATAAC
TACAGTACTGCTTCATCTCACACTGGTGGTCCACCTCAAACTTCTGCCTTTACTGAGTCGTTTGCCAGTTC
AGTAGCTGGATATGGAGACATTTACCTCAATGCTGGTCCCATGTTGTTTAGTGGAGACAAGGACCATAG
ATTTGAATATAGCCATGGTCCTATCACAGTCCTCACAAACAGCAATGACCCTTCCACAGGGCCAGAGAGT
ACTCATCCCTTGCCAGCAAAGATGCACAACTATAACTATGGTGGTAACTTACAGGAAAATCCAGGTGGC
CCTAGCCTCATGCATGGACAGACCTGGACTTCTCCTGCCCAAGGACCTGGATATTCACAAGGATACAGG
GGACACATTAGCACATCAGCTGGCAGAGGTCGAGGCAGAGGGTTACCATACTGACTCGAG

Kozak +ATG Start Codon
Flag Epitope Tag
HA Epitope Tag
Genewiz Codon Optimized Fragement
Remaining Cdk13 cDNA
Stop Codon
